# The anaerobic fungus *Caecomyces churrovis* produces H_2_ via a non-bifurcating NADH-dependent enzyme complex

**DOI:** 10.64898/2026.04.05.716614

**Authors:** Bo Zhang, Ivan Hrdy, Jan Tachezy, Yuqian Gao, Sarah M. Williams, James M. Fulcher, Nathalie Munoz, Meagan Burnet, Scott E. Baker, Michelle A. O’Malley

## Abstract

Hydrogenosomes are mitochondria-derived organelles that produce ATP and H_2_ to support energy metabolism in anaerobic eukaryotes. H_2_ production allows reoxidation of reduced cofactors generated during fermentative metabolism; however, the metabolic mechanisms for H_2_ production in anaerobic eukaryotes remains incompletely understood. In particular, it remains unclear whether anaerobic fungi (AF) hydrogenosomes use a ferredoxin-dependent pathway or a distinct mechanism to regenerate NAD(P)^+^ and link electron transfer to H_2_ formation. Here, by combining genomic search, proteomic analysis, and enzymology, we reveal the molecular mechanism for H_2_ production in the AF strain *Caecomyces churrovis*. Our enzyme assays on the organelle fraction of *C. churrovis* revealed the activity of H_2_:NAD^+^ oxidoreductase but not pyruvate:ferredoxin oxidoreductase, which is usually linked to H_2_ formation. We identified genes encoding [FeFe] hydrogenase (Hyd) and NADH dehydrogenase subunits E and F (NuoE, NuoF) in *C. churrovis*, and confirmed their expression in the isolated hydrogenosomal fractions by proteomic analysis. Combining the individually purified enzymes, we found Hyd and NuoEF proteins formed H_2_ directly from NADH independently of ferredoxin, functioning as a non-bifurcating NADH-dependent enzyme rather than an electron-bifurcating enzyme. We identified homologs of hydrogenosomal NuoE, NuoF, and Hyd in many other AF, indicating this pathway is commonly shared among the AF. This work demonstrates the existence of a non-bifurcating NADH-dependent enzyme complex in eukaryotes. Moreover, this complex could potentially be exploited as a target for controlling AF H_2_ production and altering fungal metabolism.

**IMPORTANCE:** H_2_ production is a prominent feature of anaerobic energy metabolism, yet our understanding of eukaryotic mechanisms remains limited. Anaerobic fungi (AF) are key decomposers of lignocellulose and contribute to hydrogen flux in anaerobic environments. Although it has been more than 40 years since the H_2_ production from AF was first reported, the molecular mechanism for hydrogenosomal H_2_ production and redox balance remains unclear. We demonstrate that AF produce H_2_ from NADH utilizing a non-bifurcating NADH-dependent enzyme complex rather than an electron-bifurcating, ferredoxin-dependent variant. We show that this enzyme complex is conserved across multiple AF lineages and thus demonstrate the occurrence of a non-bifurcating NADH-dependent enzyme in eukaryotes. This discovery expands our understanding of eukaryotic hydrogenosomal metabolism, reveals a previously unknown strategy for redox balancing, and highlights potential targets for manipulating H_2_ production. These insights have broad implications for microbial energy metabolism, anaerobic ecosystems, and bioengineering of H_2_-producing systems.

## INTRODUCTION

Many eukaryotes live in oxygen-limited environments where aerobic respiration is not possible. These organisms usually possess mitochondria-derived organelles, some of which support anaerobic energy metabolism. These organelles include hydrogenosomes and mitosomes, which evolved from mitochondria but differ in metabolic functions (1, 2). Hydrogenosomes have been identified in diverse anaerobic eukaryotes, including trichomonads, ciliates, and anaerobic fungi (3–5). Hydrogenosomes generate ATP through substrate-level phosphorylation and release molecular hydrogen (H_2_) as a metabolic end product (6–8). H_2_ production allows the reoxidation of reduced cofactors generated during fermentative metabolism and therefore plays an important role in maintaining redox balance in anaerobic eukaryotic cells (1, 9). Despite the importance of H_2_ production for anaerobic energy metabolism, the molecular pathways that couple electron transfer to H_2_ formation in hydrogenosomes remain incompletely understood.

H_2_ production in hydrogenosomes is typically linked to the oxidation of reduced electron carriers generated during fermentative metabolism. In trichomonads, electrons are transferred from pyruvate to ferredoxin via pyruvate:ferredoxin oxidoreductase (PFOR), and reduced ferredoxin donates electrons to an [FeFe] hydrogenase to produce H_2_ (10). In an anaerobic ciliate *Nyctotherus ovalis*, it was proposed that NADH donates electrons to form H_2_ via a monomeric multi-domain [FeFe] hydrogenase (a fusion of [FeFe] hydrogenase and NADH dehydrogenase subunit E and F) (11, 12), but its biochemical evidence is missing. Anaerobic fungi (AF) also encode [FeFe] hydrogenases and other anaerobe-specific enzymes (13, 14), and their hydrogenosomes generate ATP and H_2_ (15, 16), suggesting a central role in energy metabolism. AF are key members of the herbivore gut microbiome, where they degrade lignocellulose and release H_2_ that supports syntrophic interactions with methanogens and other microorganisms (17, 18). They hold potential for biomass conversion and bioproduct formation in anaerobic microbial consortia (19–21), including medium-chain fatty acids (22, 23) and hydrogen-based bioenergy (24). AF are a potential source of biological H_2_ in the oceanic crust, and may be involved in fundamental biogeochemical cycles by providing H_2_ to prokaryotes in the deep biosphere (25, 26). Despite this importance and decades of study, it remains unclear whether AF hydrogenosomes use a ferredoxin-dependent pathway or a distinct NAD(P)^+^-dependent mechanism of H_2_ formation (Figure 1A) (13, 27, 28), making it difficult to design accurate genome-scale metabolic models of AF usable in biotechnological and industrial applications (13).

**Figure 1.**
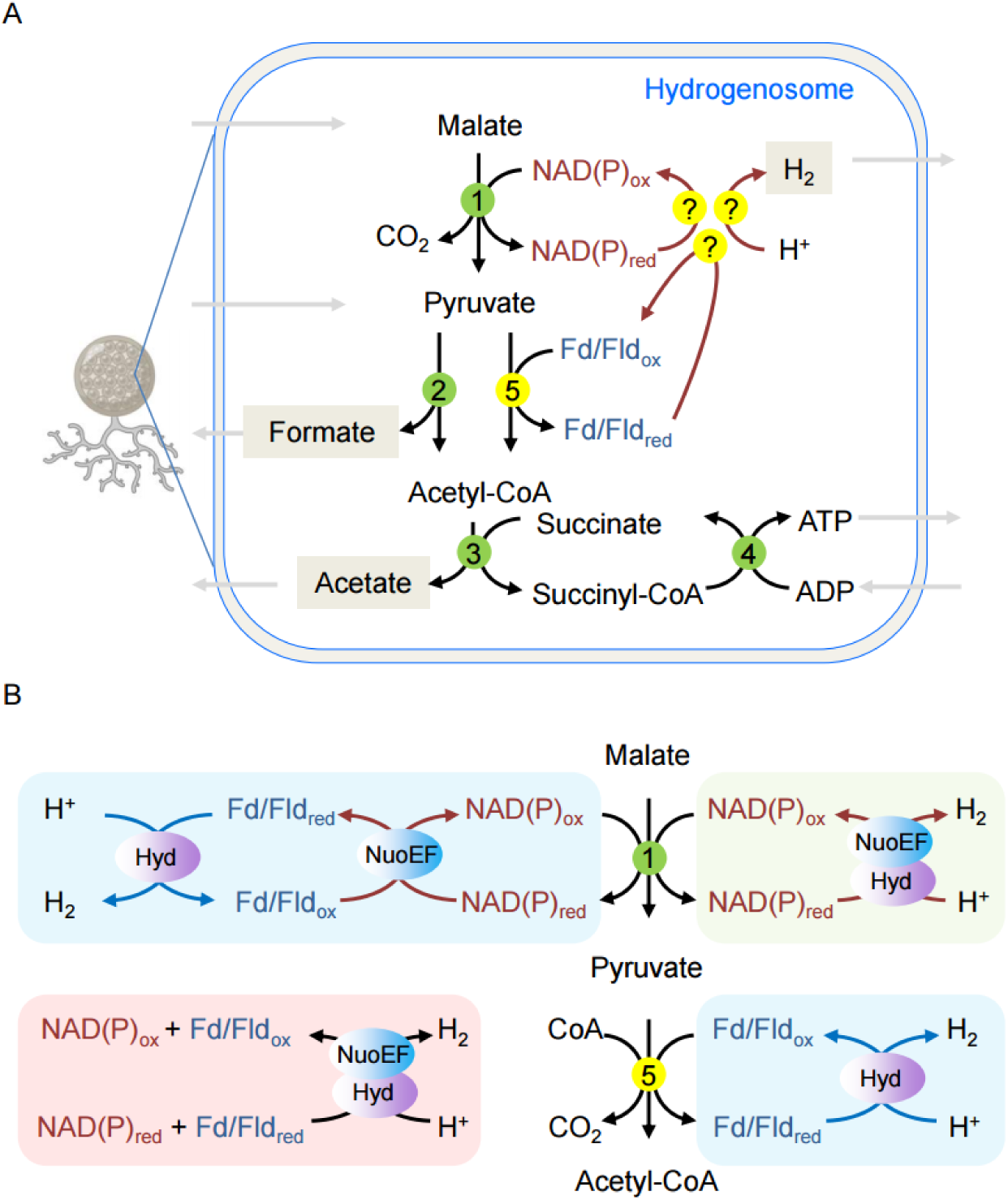
H_2_ production pathways in the hydrogenosomes of anaerobic gut fungi (AF) are unknown. (A) Unknown pathways for H_2_ production in anaerobic gut fungi; (B) Possible enzyme pathways for forming H_2_ in the hydrogenosome of anaerobic gut fungi. NAD(P)_ox_, NAD(P)^+^; NAD(P)_red_, NAD(P)H; Fd/Fld_ox_, oxidized ferredoxin or flavodoxin; Fd/Fld_red_, reduced ferredoxin or flavodoxin; 1, malic enzyme; 2, pyruvate formate lyase; 3, acetyl-CoA:succinate CoA transferase; 4, succinyl-CoA synthetase; 5, possible pyruvate:ferredoxin/flavodoxin oxidoreductase; Hyd, [FeFe] hydrogenase; NuoEF, NADH dehydrogenase (complex I) subunits E and F; “?” for unknown pathways for recycling NAD(P) and/or ferredoxin/flavodoxin and forming H_2_.

Recent genomic analysis of AF identified genes encoding an [FeFe] hydrogenase and the NADH dehydrogenase (mitochondrial respiratory complex I) subunit E and F (NuoEF) (13, 14).These enzymes could contribute to H_2_ production and redox balance through several possible mechanisms (Figure 1B) (1, 9, 10, 13, 14), depending on whether NAD(P)H and/or ferredoxin act as electron donors. First, ferredoxin is reduced by electrons generated from pyruvate by the activity of pyruvate:ferredoxin oxidoreductase (PFOR). Alternatively, electrons are released from malate by malic enzyme activity, which utilizes NAD(P)^+^ as electron acceptor, and transferred to ferredoxin by NAD(P)H:ferredoxin oxidoreductase (complex I). The reduced ferredoxin then provides electrons for H_2_ production by the [FeFe] hydrogenase, as observed in trichomonads hydrogenosomes (9). When NAD(P)H is the only electron donor, hydrogenase could couple with NuoEF to form H_2_ directly from NAD(P)H, as suggested by several research groups (14, 29, 30). These hydrogenases are called non-bifurcating NAD(P)H-dependent hydrogenase as they catalyze the direct, reversible oxidation of NAD(P)H to produce H_2_ (or vice versa) without the need for electron bifurcation or reduced ferredoxin (31). <u>This</u> mechanism was initially proposed for the *N. ovalis* monomeric [FeFe] hydrogenase (11, 12), but its biochemical evidence is missing. Lastly, both NAD(P)H and reduced ferredoxin could donate electrons for H_2_ production by the hydrogenase and NuoEF using the bifurcating/confurcating mechanism, as speculated to occur in hydrogenosomes of *Trichomonas vaginalis* (1, 10) and shown in an analogous case of bacterial trimeric hydrogenase, e.g. in *Thermotoga maritima* (32). It is under debate whether a PFOR is involved in H_2_ synthesis in AF hydrogenosomes (8). Although the activity of PFOR was observed in *Neocallimastix* sp. L2 and *Neocallimastix patriciarum* (5, 16), it was undetectable in *Neocallimastix frontails* (33). Pyruvate is predominantly metabolized by pyruvate formate lyase (PFL) in AF, which does not release electrons for H_2_ synthesis (29). Thus, it remains unclear how H_2_ is generated in AF hydrogenosomes.

Here, we elucidated the pathway responsible for H_2_ production in the hydrogenosome of the model anaerobic fungus *Caecomyces churrovis* from a sheep digestive tract. By the combination of metabolic phenotyping, comparative genomics, proteomic analysis, protein purification, and reconstituted enzyme assays, we demonstrated that the Hyd and NuoEF of *C. churrovis* form H_2_ from NADH independently of ferredoxin, functioning as a non-bifurcating NADH-dependent enzyme complex which is apparently commonly present in many anaerobic fungi. These results define a previously unrecognized, non-bifurcating NADH-dependent hydrogenase pathway in eukaryotic microorganisms and provide a framework for understanding how anaerobic fungi couple electron transfer to H_2_ production. This work also highlights potential targets to manipulate AF metabolism for applications in anaerobic ecosystems and bioenergy production.

## RESULTS

### *C. churrovis* forms H_2_ using H_2_:NAD^+^ oxidoreductase pathway

To confirm that *C. churrovis* forms H_2_ during glucose fermentation, we measured H_2_ released to the headspace of a cultivation bottle under anaerobic conditions. We found that *C. churrovis* produced about 106 µmol H_2_ from 767 µmol glucose after 7 days of growth (Figure 2, Supplementary table 1). In addition to H_2_, other major fermentation products were detected, including formate, ethanol, acetate, lactate, and succinate (Supplementary table 1).

**Figure 2.**
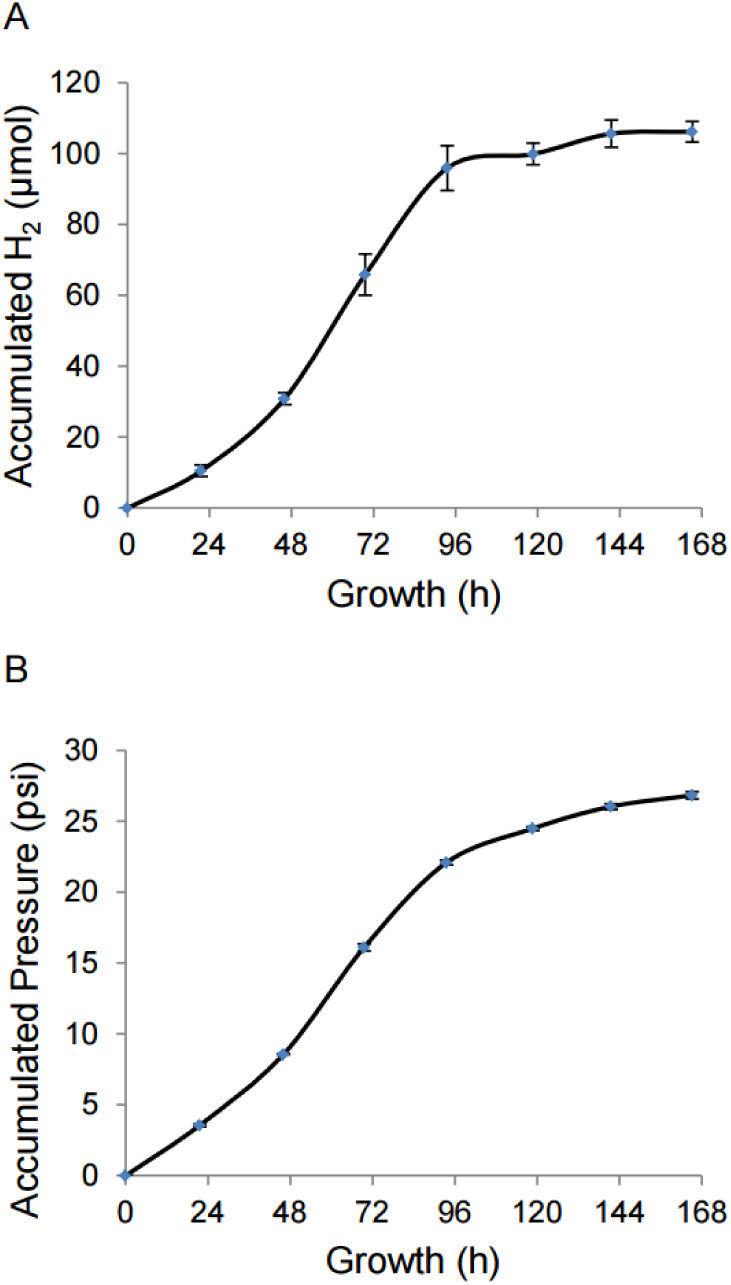
Accumulated H_2_ production and pressure by *C. churrovis* in 40 mL medium MC minus with 5 g/L glucose. H_2_ content was measured on a daily basis for *C. churrovis* as described in the methods. Briefly, 2 mL 3-days age seed was inoculated to 39 mL medium MC minus with glucose in a 70-mL serum bottle, from which 1 mL culture was taken for 0 h point fermentation products analysis. The remaining 40 mL culture was incubated at 39°C. Gas pressure was measured and released to 0 psi monitored by a pressure meter on a daily basis. Four cultures were included as biological replicates.

To identify which pathway is used for H_2_ formation in the anaerobic gut fungus *C. churrovis* (Figure 1B), we measured the enzyme activities in the large organelle fraction (LOF) of *C. churrovis* (Table 1). First, we measured the activity of malic enzyme, a marker enzyme of hydrogenosomes (16). High enzyme activities of NAD(P)^+^-dependent malic enzyme were observed in the LOF, confirming the presence of hydrogenosomes in the LOF. Subsequently, we measured the reduction of NAD^+^ and ferredoxin with H_2_ in the LOF. If H_2_ is formed using electrons from NADH, the reduction of NAD^+^ with H_2_ by the LOF should occur. If H_2_ is formed using electrons from reduced ferredoxin only (not NADH), we should detect the reduction of ferredoxin with H_2_ by the LOF. We detected H_2_:NAD^+^ oxidoreductase activity (mean ± SEM, 80 ± 27 mU/mg) in the LOF (Table 1, Supplementary Figure 1). The addition of a heterologously generated ferredoxin of *C. churrovis* (CcFd-Strep, referred as “CcFd” in the following text) to the assay did not increase the reduction rate of NAD^+^ with H_2_, and no reduction of CcFd with H_2_ was observed (Supplementary Figure 1). This suggests that H_2_ formation is catalyzed by the H_2_:NAD^+^ oxidoreductase pathway using NADH as a sole electron donor. In addition, we observed hydrogenase activity [H_2_:methyl viologen (MV) oxidoreductase] and NADH dehydrogenase (NADH:MV oxidoreductase) activity in the LOF, demonstrating the existence of hydrogenase and NADH dehydrogenase (complex I) in the LOF.

**Table 1.** Enzyme activity measured in the large organelle fraction (LOF) of *C. churrovis*.

| Enzyme activity | Mean ± SEM<br>(mU/mg protein) | *P*-value |
| --- | --- | --- |
| Malic enzyme (NAD<sup>+</sup>) | 1522 ± 432 | 0.0360 |
| Malic enzyme (NADP<sup>+</sup>) | 732 ± 190 | 0.0308 |
| H<sub>2</sub>:NAD<sup>+</sup> oxidoreductase (no ferredoxin) | 80 ± 27 | 0.0499 |
| H<sub>2</sub>:NAD<sup>+</sup> oxidoreductase (before CcFd addition) | 18 ± 4 | 0.0225 |
| H<sub>2</sub>:NAD<sup>+</sup> oxidoreductase (after CcFd addition) | 18 ± 3 | 0.0155 |
| H<sub>2</sub>:NADP<sup>+</sup> oxidoreductase | Not detectable | Not applicable |
| H<sub>2</sub>:MV oxidoreductase | 20 ± 11 | 0.1065 |
| NADH:MV oxidoreductase | 399 ± 92 | 0.0245 |
| Pyruvate:MV oxidoreductase | Not detectable | Not applicable |
| Pyruvate:CcFd oxidoreductase | Not detectable | Not applicable |

To test if PFOR is involved in the reduction of ferredoxin, we measured its activity using MV or CcFd as an electron acceptor, respectively. We did not observe any pyruvate:MV oxidoreductase or pyruvate:CcFd oxidoreductase activity in the LOF. Additionally, we spiked the cytoplasmic fraction of *Prevotella brevis* GA33, a positive control for such activity (34), at the end of the experiments and we observed activity of pyruvate:MV oxidoreductase (83 ±17 mU/mg) and pyruvate:CcFd oxidoreductase (139 ±13 mU/mg) (Supplementary Figure 1). This supports that our assay conditions were appropriate to detect the PFOR activity.

Next, we looked for genomic evidence of observed enzymatic activities and searched for hydrogenase and NADH dehydrogenase genes in the genome of *C. churrovis* (35). We found the genes encoding [FeFe] hydrogenase (Hyd, protein ID 243125 in MycoCosm), NADH dehydrogenase subunit E (NuoE, 24-kDa subunit, protein ID 454874), and NADH dehydrogenase subunit F (NuoF, 51-kDa subunit, protein ID 557447). DeepLoc2 predicted that all three proteins possess an N-terminal targeting signal for hydrogenosomal localization, suggesting that they are hydrogenosomal proteins. Unlike some other proteins (e.g. pyruvate formate lyase) that have several paralogs in the genome, we did not find other paralogs of Hyd, NuoE, and NuoF in the genome of *C. churrovis*.

### Proteomic analysis verified expression of key hydrogenosomal proteins in AF

To verify the expression of Hyd, NuoE, and NuoF, we performed nanoPOTS proteomic analysis on the enriched hydrogenosomal fractions. First, we established a method to isolate enriched hydrogenosomes from *C. churrovis*. Briefly, we lysed the cells, prepared crude organelle fractions by differential centrifugation, and separated organelles by OptiPrep density gradient centrifugation (Figure 3A). We collected eight fractions and measured the activity of malic enzyme (NADP^+^), aiming to identify the fraction(s) that contained enriched hydrogenosomes. We found three fractions (#4-6) with the highest malic enzyme (NADP^+^) activity (Figure 3B). We further confirmed the presence of hydrogenosomes in these fractions by western blot using the anti-serum against the β subunit of succinyl-CoA synthetase (βSCS), which is a hydrogenosomal marker protein in AF (36) (Supplementary Figure 2).

**Figure 3.**
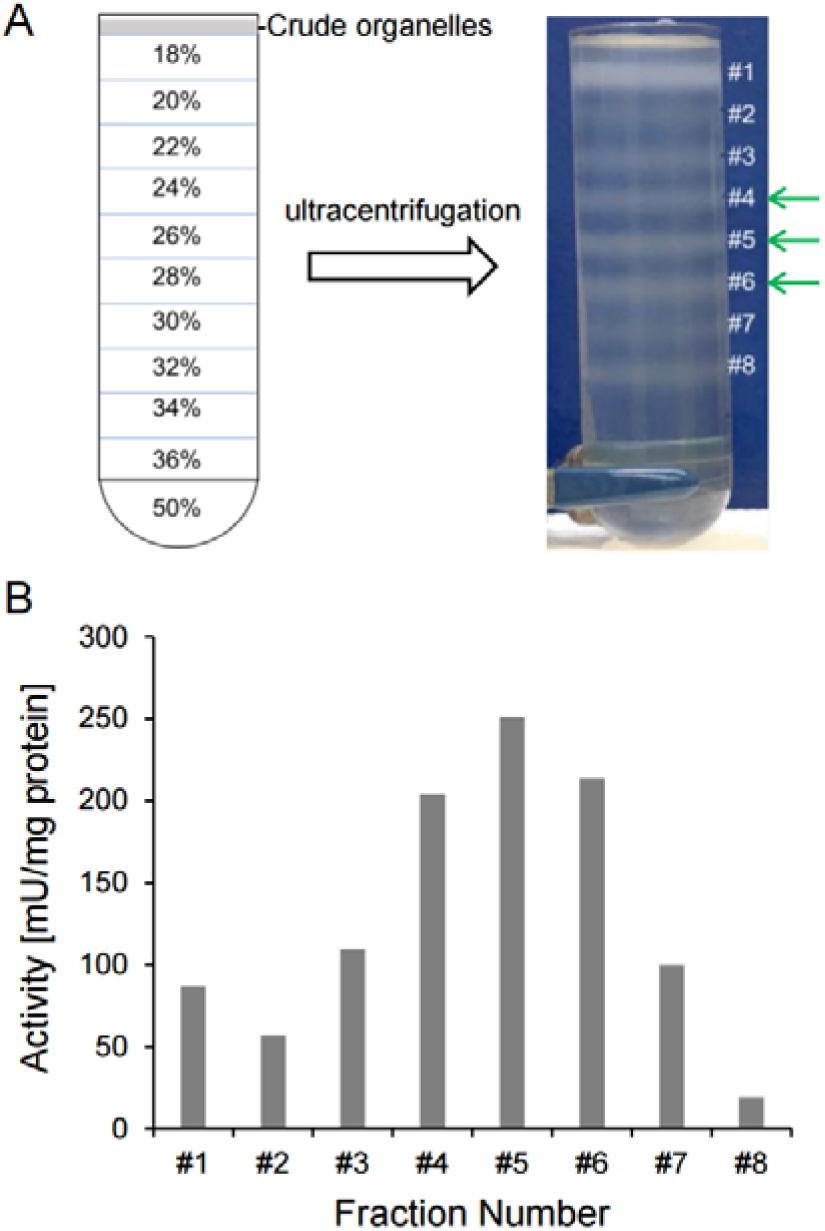
Hydrogenosome fractions from *C. churrovis* were isolated via OptiPrep density gradient and identified by enzyme assays. (A) Isolation of hydrogenosomes by OptiPrep density gradient centrifugation. (B) Fraction #4, 5, 6 has the highest malic enzyme (NADP^+^) activity. Enzyme activity shown here represents one of three independently prepared fractions.

Using this method we prepared triplicates of enriched AF hydrogenosome fractions (#4-6) for nanoPOTS proteomic analysis, which allows deep and quantitative proteomic profiling of low abundance samples (37). Both NADP^+^ and NAD^+^-dependent malic enzyme activities were observed in fraction #4-6, with the NAD^+^-dependent activity being approximately 3 fold of the NADP^+^-dependent activity in all three fractions (Supplementary Table 2). Our nanoPOTS proteomic analysis of the enriched hydrogenosomes confirmed expression of several key enzymes, including Hyd, NuoE, and NuoF (Table 2, Supplementary Figure 3). Specifically, reproducible MS/MS spectra of numerous unique peptides corresponding to these proteins were detected across three biological replicates, demonstrating their expression in the cells under the experimental conditions (Supplementary Dataset 1). Interestingly, no MS/MS spectrum of the hydrogenosomal ferredoxin (CcFd) was detected in two biological replicates and only one MS/MS spectrum was detected in the third biological replicate with low intensity (Supplementary Figure 4). Our proteomic analysis also detected a protein annotated as pyruvate:ferredoxin/flavodoxin oxidoreductase or sulfite reductase. However, our enzyme assay did not detect any activity of PFOR, leaving the function of the PFOR homolog in the fungal hydrogenosome unknown.

**Table 2.**
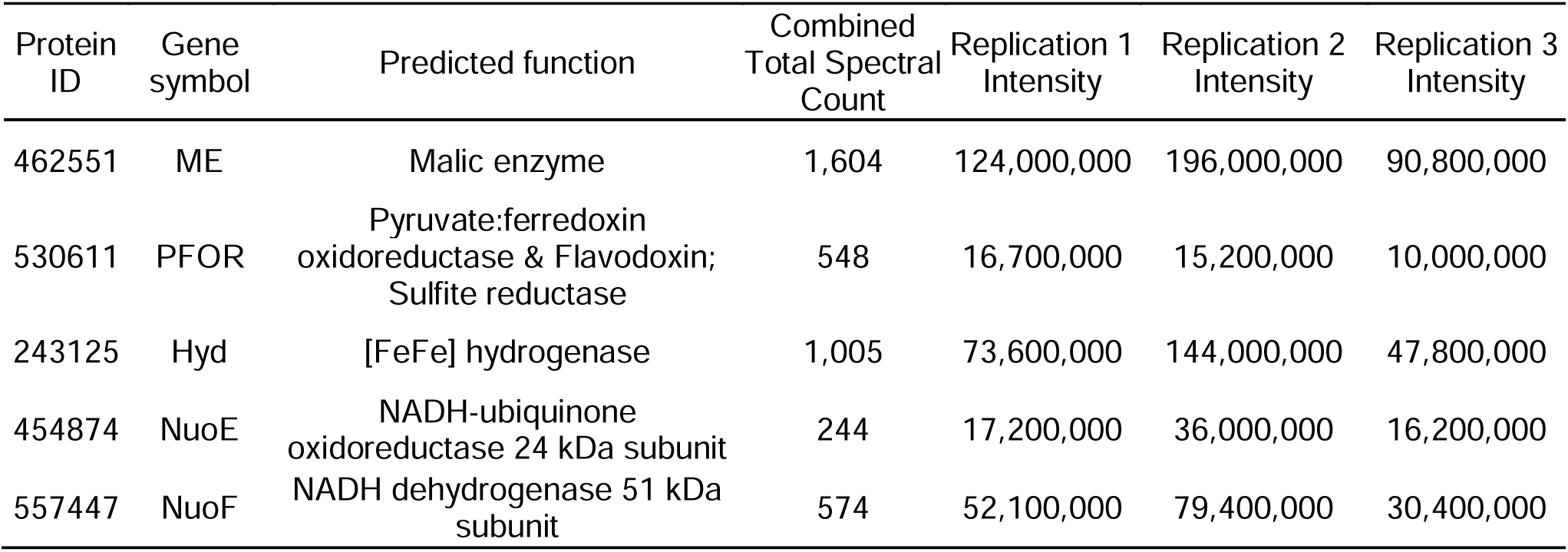
NanoPOTS proteomic analysis of the pooled hydrogenosome fractions confirmed expression of several key enzymes.

| Protein ID | Gene symbol | Predicted function | Combined Total Spectral Count | Replication 1 Intensity | Replication 2 Intensity | Replication 3 Intensity |
| --- | --- | --- | --- | --- | --- | --- |
| 462551 | ME | Malic enzyme | 1,604 | 124,000,000 | 196,000,000 | 90,800,000 |
| 530611 | PFOR | Pyruvate:ferredoxin oxidoreductase & Flavodoxin; Sulfite reductase | 548 | 16,700,000 | 15,200,000 | 10,000,000 |
| 243125 | Hyd | [FeFe] hydrogenase | 1,005 | 73,600,000 | 144,000,000 | 47,800,000 |
| 454874 | NuoE | NADH-ubiquinone oxidoreductase 24 kDa subunit | 244 | 17,200,000 | 36,000,000 | 16,200,000 |
| 557447 | NuoF | NADH dehydrogenase 51 kDa subunit | 574 | 52,100,000 | 79,400,000 | 30,400,000 |

We also performed global proteomic analysis of cell lysate of *C. churrovis*. Peptides of proteins listed in Table 2 were also detected in the cell lysate (Supplementary Table 3), and no peptides derived from CcFd were detected.

### Reconstruction of H_2_ producing pathway *in vitro*

Our enzyme assays, genomic analysis, and proteomic analysis suggest that the H_2_:NAD^+^ oxidoreductase pathway is used for H_2_ production using electrons from NADH but not from ferredoxin. To further investigate the function of this system, we prepared recombinant [2Fe2S] ferredoxin, NuoE and NuoF proteins, and hydrogenase for targeted enzyme assays.

First, we cloned the gene encoding a hydrogenosomal [2Fe2S] ferredoxin in *Trichomonas vaginalis* (TvFd, TVAG_003900 in the TrichDB database) and *C. churrovis* (CcFd, protein ID 549362 in MycoCosm). TvFd is an experimentally verified [2Fe-2S] ferredoxin and functions as an electron carrier in the hydrogenosome of *T. vaginalis* (38, 39). Matured TvFd and CcFd (TvFd and CcFd without their hydrogenosomal targeting sequences) were heterologously generated in *E. coli* BL21(DE3)Δ*iscR* and purified (Figure 4A). To test their native function, we prepared enzymes that are capable of using ferredoxin as an electron carrier. These enzymes include a recombinant 2-oxoglutarate:ferredoxin oxidoreductase (OGOR) of *Prevotella ruminicola* 23 and the crude hydrogenase of *Clostridium pasteurianum* W5 (40). We confirmed that both ferredoxins are functional. The recombinant OGOR reduced TvFd and CcFd with 2-oxoglutarate (Figure 4B). The specific activity of 2-oxoglutarate:TvFd oxidoreductase was 0.698 ± 0.234 U/mg and 2-oxoglutarate:CcFd oxidoreductase was 0.295 ± 0.064 U/mg, respectively. Similarly, we observed the reduction of TvFd and CcFd with H_2_ by the crude hydrogenase of *C. pasteurianum* W5, with the specific activity of H_2_:TvFd oxidoreductase at 111.253 ± 1.121 U/mg and H_2_:CcFd oxidoreductase at 13.807 ± 1.458 U/mg (Figure 4B). Therefore, our recombinant TvFd and CcFd can function as an electron carrier in the enzyme assays.

**Figure 4.**
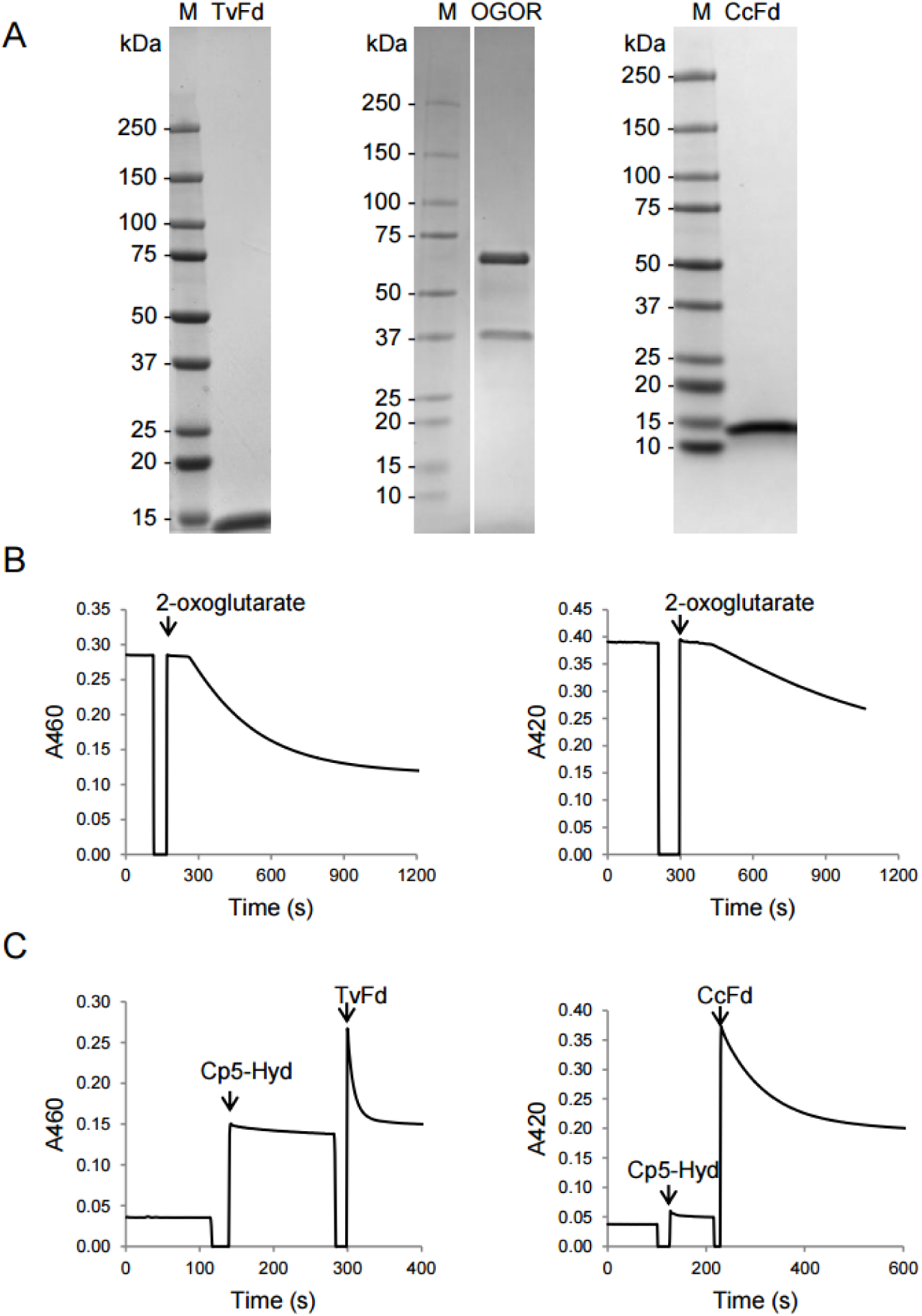
Heterologously overexpressed TvFd, CcFd, and OGOR were purified and they are functional in enzyme assays. (A) Purified proteins were analyzed by SDS-PAGE and stained with Coomassie G-250. (B) TvFd (left panel) or CcFd (right panel) was reduced by OGOR with 2-oxoglutarate. (C) TvFd (left panel) or CcFd (right panel) was reduced with H_2_ by the crude hydrogenase of *C. pasteurianum* W5 (Cp5-Hyd). In panel A, “M” is Precision Plus Protein Dual Color Standards (Bio-Rad #1610374). “TvFd” is 4 µg TvFd with N-terminal Strep tag II (11.1 kDa). “OGOR” is 3 µg recombinant OGOR, including its α subunit with Strep-tag II at N-terminal (67 kDa) and β subunit (36.6 kDa). “CcFd” is 2 µg CcFd with C-terminal Strep tag II (13.1 kDa). In panel B, the enzyme assay was initiated by the addition of TvFd or CcFd. In panel C, the assay was bubbled with H_2_ for 5 min before measuring the absorbance. See the methods for details about enzyme components. The enzyme assays shown here are one representative of two technical replicates for (B) and three technical replicates for (C).

Next, we prepared a recombinant NuoE-NuoF complex of *C. churrovis*. NuoE and NuoF were co-expressed in the plasmid pET32a (see method section for plasmid construction). We purified the NuoE together with NuoF using the C-terminal Strep-tag II of NuoF. The co-purification of NuoE and NuoF demonstrates that they formed a complex (NuoEF-Strep) (Figure 5A). The purified NuoEF-Strep displayed high enzymatic activity of NADH:MV oxidoreductase (26.452 ± 7.490 U/mg) (Figure 5C). We tested if other electron carriers can be reduced with NADH, and we found that NuoEF was not able to reduce CcFd or TvFd (Supplementary Figure 5).

**Figure 5.**
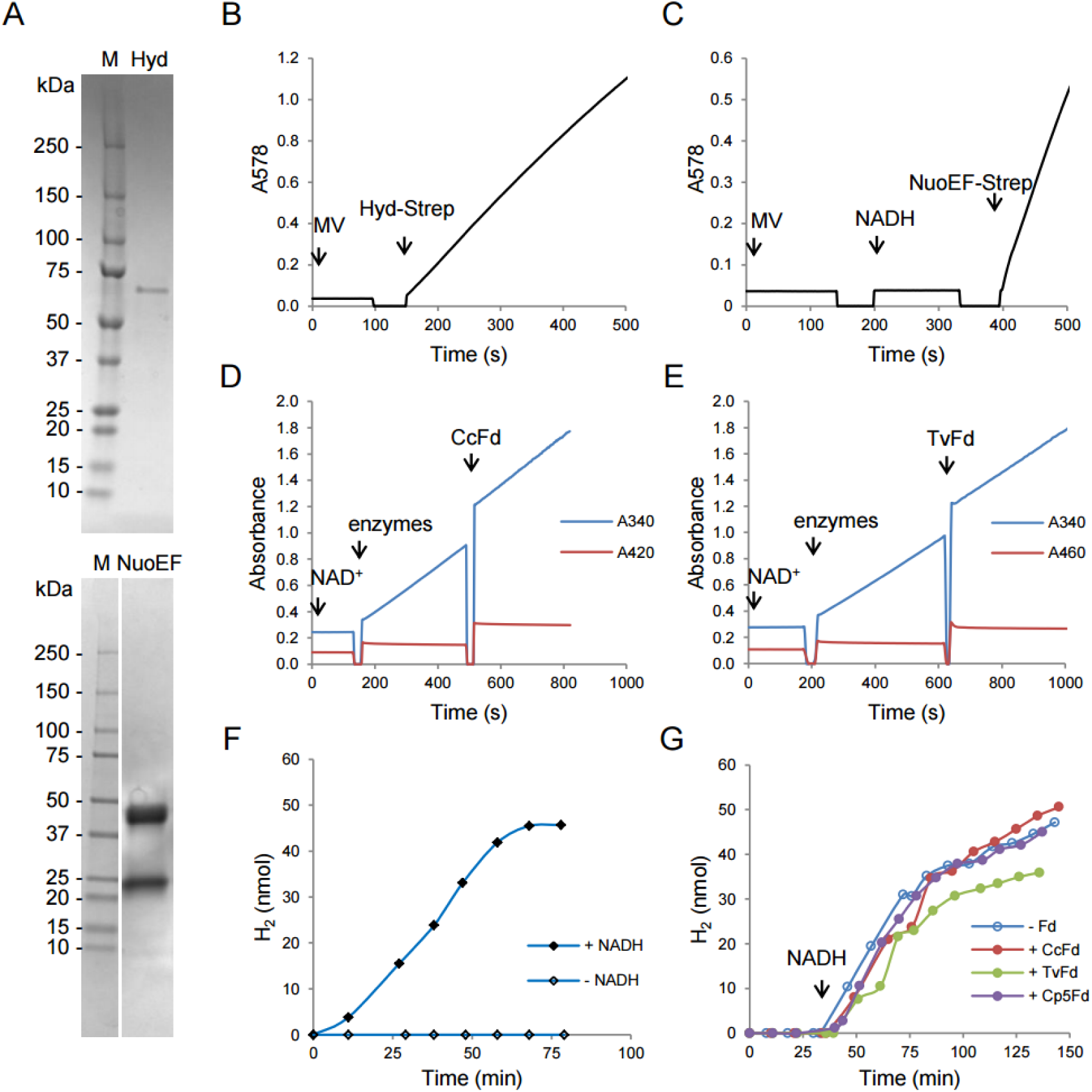
Hyd-Strep and NuoEF-Strep function as a non-bifurcating NADH-dependent enzyme. (A) Purified protein (1.3 µg Hyd-Strep or 3.6 µg NuoEF-Strep) was analyzed via SDS-PAGE and stained with Coomassie G-250. “M” is Precision Plus Protein Dual Color Standards (Bio-Rad #1610374). Hyd-Strep (in lane “Hyd”) is 69.6 kDa, NuoE (in lane “NuoEF”) is 23.6 kDa, and NuoF-Strep is 49.6 kDa. (B) Methyl viologen (MV) was reduced by Hyd-Strep, showing the detectable H_2_:MV oxidoreductase activity of the purified Hyd-Strep. The assay containing MV was bubbled with H_2_ for 5-10 min before measuring absorbance at 578 nm. (C) MV was reduced by NADH after addition of NuoEF-Strep, showing the detectable NADH:MV oxidoreductase activity of the NuoEF-Strep. (D-E) The mixture of Hyd-Strep and NuoEF-Strep reduced NAD^+^ with H_2_ independent of CcFd (D) and TvFd (E), but didn’t reduce CcFd and TvFd with H_2_. The assay containing 1 mM NAD^+^ was bubbled with H_2_ for 5-10 min before measuring its absorbance. The enzymes (mixture of Hyd-Strep and NuoEF-Strep) and 30 µM CcFd or TvFd were then added. (F) The mixture of Hyd-Strep and NuoEF-Strep catalyzed H_2_ formation directly from NADH. Hyd-Strep was mixed with NuoEF-Strep under N_2_ atmosphere at room temperature for 5 min before being added to the assay. NADH (2 mM) was added to the assay to start the reaction and water was used as the negative control (“- NADH”). Then a gas sample was taken for H_2_ detection by gas chromatography. (G) H_2_ was not formed from reduced ferredoxin but from NADH. “+ CcFd”, “+ TvFd”, and “+ Cp5Fd” represent the reactions containing reduced CcFd, TvFd, or the *C. pasteurianum* W5 ferredoxin. “- Fd” represents that no any ferredoxin was added. All these reactions contained the Fd-reducing system (2-ketoglutarate, CoA, TPP, MgCl_2_, and OGOR) and were incubated at 39°C for 30 min to allow reduction of ferredoxin, if ferredoxin was added. Hyd-Strep and NuoEF-Strep were mixed for 5 min at room temperature and then added to the assay to start the reaction under N_2_. Gas samples were taken over time for H_2_ detection by gas chromatography. NADH (2 mM) was added to the assay after about 30-35 min since adding enzyme mixture. The maximum rates of forming H_2_ after adding NADH were at the same level (no statistically significant difference) in the four reactions. One representative figure for each assay is shown here. Two batches of independently purified samples (Hyd-Strep, NuoEF-Strep) were tested. See the method section for details about enzyme assays.

Hydrogenase of *C. churrovis* was coexpressed with the hydrogenase maturases of *Shewanella oneidensis* MR-1 as hydrogenase maturases are required to assemble the H-cluster of [FeFe] hydrogenase. The affinity-purified hydrogenase (C-terminally tagged, Hyd-Strep) revealed high specific activity of H_2_:MV oxidoreductase (5.607 ±1.882 U/mg), (Figure 5B). We then tested if other electron carriers can be reduced with H_2_. We found that this hydrogenase was not able to use H_2_ for the reduction of NAD^+^, NADP^+^, CcFd, and TvFd (Supplementary Figure 5). The purity of recombinant NuoEF complex and hydrogenase was confirmed by LC-MS/MS analysis (Supplementary Figure 6). None of the contaminating proteins in the Hyd-Strep and NuoEF-Strep samples from *E. coli* was predicted to have hydrogenase or NADH dehydrogenase activity (Supplementary Dataset 2 and 3).

We then tested if the mixture of *C. churrovis* Hyd-Strep and NuoEF-Strep can use H_2_ to reduce NAD^+^ or ferredoxin or both. We detected a reduction of NAD^+^ to NADH in the presence of H_2_ after addition of the enzyme mixture to the assay in the absence of ferredoxin (Figure 5D, 5E). The measured specific activity of NAD^+^ reduction with H_2_ by the enzyme mixture varies from 0.840 U/mg to 3.451 U/mg, depending on the sample batches. Addition of ferredoxin (CcFd or TvFd) to the assays did not increase the NAD^+^ reduction rate, showing that the enzyme mixture can reduce NAD^+^ using H_2_ independently of ferredoxin. In addition, no reduction of CcFd or TvFd with H_2_ was observed in the presence of NAD^+^ (Figure 5D, 5E), showing that ferredoxin is not involved in H_2_ consumption in this enzyme assay. In sum, we found that the enzyme mixture of Hyd-Strep and NuoEF-Strep can reduce NAD^+^ using H_2_ as a source of electrons independently of ferredoxin, and the combined enzymes cannot use H_2_ to reduce ferredoxin.

Next, we tested if the mixture of Hyd-Strep and NuoEF-Strep can use electrons from NADH to form H_2_. After adding the mixture of both enzymes into the assay containing NADH, we observed H_2_ production at the rate of 98 ± 1 nmol H_2_/min/mg protein (Figure 5F, Supplementary Table 4). In comparison, no H_2_ was formed in the absence of NADH (Figure 5F), demonstrating that NADH can contribute directly to H_2_ production by the enzyme mixture. The activity was strictly NADH-specific as no H_2_ production was observed using NADPH as an electron source (Supplementary Figure 7). Then we investigated whether reduced ferredoxins (CcFd or TvFd) can provide electrons for H_2_ production instead of NADH. Ferredoxin was reduced with 2-oxoglutarate by the recombinant OGOR in the presence of coenzyme A (CoA). We did not observe H_2_ production from the reduced ferredoxin after adding the mixture of Hyd-Strep and NuoEF-Strep to the assay until NADH was added. The maximum rate of H_2_ formation from NADH in the absence of ferredoxin was not statistically different from the assays containing reduced ferredoxin and NADH, demonstrating that the reduced ferredoxin did not contribute to H_2_ production by the enzyme mixture (Figure 5F, 5G, Supplementary Table 4). In the control assay, we confirmed that reduced ferredoxin is functional and can serve as an electron carrier for H_2_ production by “standard” hydrogenase, since we observed H_2_ formation after adding the crude hydrogenase of *C. pasteurianum* W5 to the ferredoxin-reducing system (H_2_ production at the rate of 32 ± 1, 36 ± 13, 55 ± 17 nmol H_2_/min/mg protein from reduced CcFd, TvFd, and ferredoxin of *C. pasteurianum* W5, respectively).

In summary, our enzyme assays with *C. churrovis* Hyd-Strep and NuoEF-Strep demonstrate that these enzymes have NADH-dependent non-bifurcating enzyme activity, but not bifurcating hydrogenase activity nor ferredoxin-dependent hydrogenase activity. They are involved in H_2_ production using NADH directly as electron donor, but not NADPH or ferredoxin.

### Hydrogenase, NuoE, and NuoF interact to form a complex

The structure prediction of protein interactions by AlphaFold Server (41) between *C. churrovis* hydrogenosomal hydrogenase, NuoE, and NuoF suggested that these proteins may form a complex similar to the predicted structure of Hyd1ABC of *Syntrophomonas wolfei* (Supplementary Figure 8) (42). Thus, we tested experimentally whether the complex is formed upon co-expression of all three components in *E. coli* BL21(DE3)Δ*iscR.* N-terminally tagged hydrogenase (Strep-Hyd) was used as a bait to purify the complex. Indeed, NuoE and NuoF were co-eluted from the resin together with hydrogenase, as shown in three eluted fractions (B2, B3, B4) visualized by SDS-PAGE (Figure 6B). Our native PAGE analysis of the purified proteins (eluted fractions) identified two protein complex bands, and they disappeared in the sample denatured by boiling before the electrophoresis (Figure 6C). The presence of corresponding proteins in each of the two gel bands was confirmed by LC-MS/MS analysis, demonstrating that hydrogenase, NuoE, and NuoF form a complex (Supplementary Figure 9, Supplementary Dataset 4 and 5). Moreover, we observed high NADH dehydrogenase activity (NADH:MV oxidoreductase) in the elution fractions B2 (176.977 ± 5.144 U/mg), B3 (151.307 ± 4.962 U/mg), and B4 (168.531 ± 5.026 U/mg), confirming the existence of NuoEF in the elution fractions. However, we did not detect H_2_:MV oxidoreductase activity, although Strep-Hyd was coexpressed with the *S. oneidensis* hydrogenase maturases together with NuoE, and NuoF in the same *E. coli* cell.

**Figure 6.**
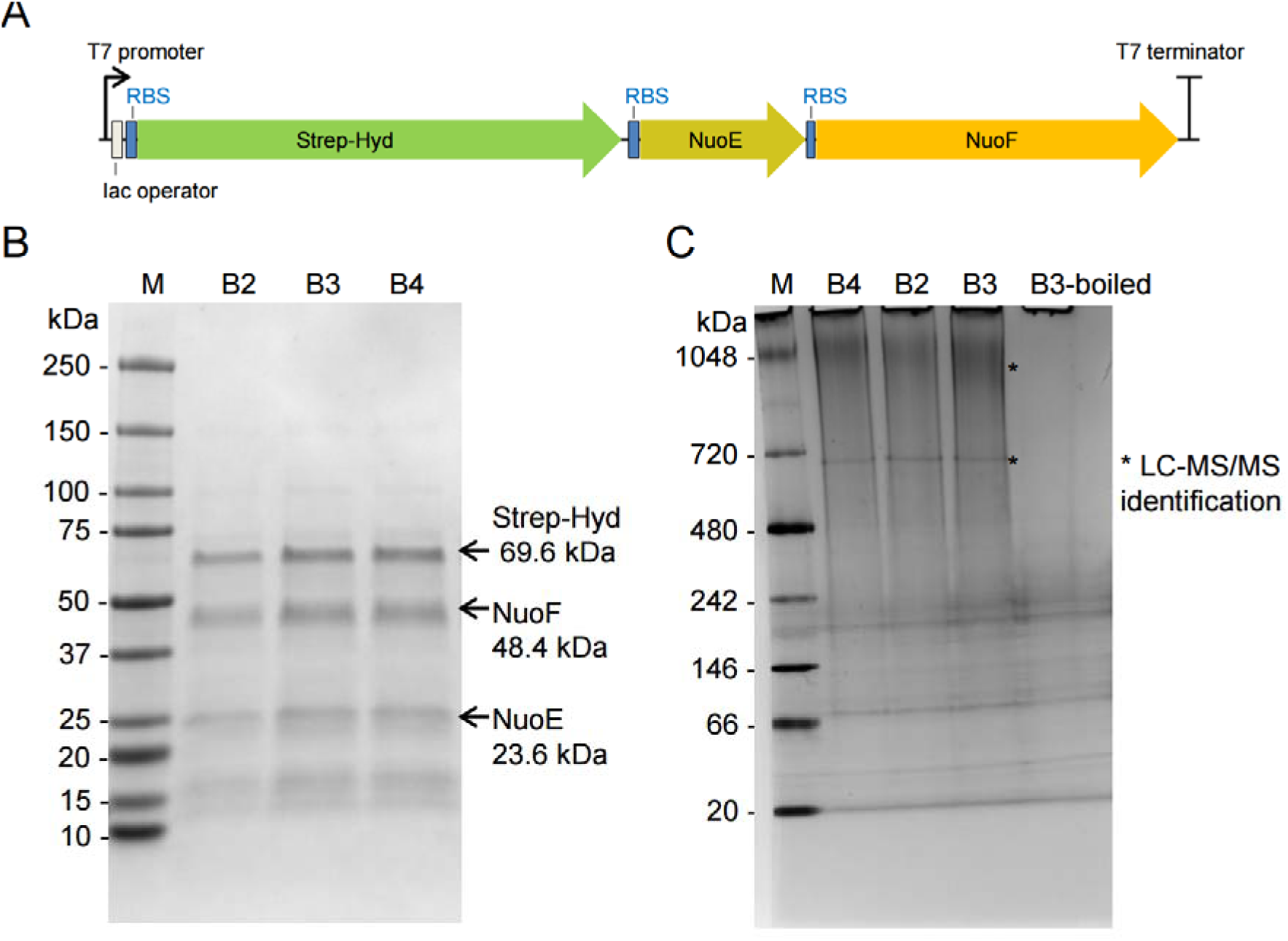
Co-purification of Hyd, NuoE, and NuoF demonstrates that they interact to form a complex. (A) A diagram showing the order of Strep-Hyd, NuoE, and NuoF located between T7 promoter and T7 terminator in pET32a. Strep-tag II was located at the N-terminal of Hyd, but no tag was added to NuoE and NuoF. RBS, ribosome binding site from bacteriophage T7 gene. The *E. coli* cell was transformed with both pET32a-Strep-Hyd-NuoEF and pACYCDuet1-HydGX-HydEF. (B) Three elution fractions (B2-B4) purified by Strep-Tactin Superflow resin were analyzed by SDS-PAGE and stained with Coomassie G-250. (C) The elution fractions were analyzed by Native-PAGE (4-16%) and stained with coomassie-G-250. Lane B2, B3, B4 are for samples (0.89, 1.31, 1.37 µg for B2-B4) in native status (without denaturation). Lane B3-boiled is for B3 fraction that was denatured by boiling. Two gel bands (smearing and sharp band) indicated by the asterisk symbol were cut. Strep-Hyd, NuoE, NuoF were detected in both bands by LC-MS/MS.

### Hydrogenosomal Hyd, NuoE, and NuoF are conserved in AF

Having established that the hydrogenosomal Hyd, NuoE, and NuoF are responsible for H_2_ production in *C. churrovis*, we investigated whether other AF can potentially use the same pathway for H_2_ production in hydrogenosomes. We searched for homologs of these proteins in 12 genomes of AF with publicly available genome information by July 2025. These AF belong to five genera (*Anaeromyces*, *Caecomyces, Neocallimastix, Pecoramyces,* and *Piromyces*) across four families of Neocallimastigales order. We found the homologs of Hyd, NuoE and NuoF in 11 fungi (Table 3). Interestingly, we identified two homologs of Hyd, NuoE, and NuoF in the genus *Neocallimastix*, while only one homolog of each protein in the genera *Anaeromyces, Caecomyces, Pecoramyces,* and *Piromyces.* Their identities with Hyd, NuoE, and NuoF of *C. churrovis* are in the range of 86-91%, 90-92%, and 90-93%, respectively. In the *Piromyces sp.* E2, NuoE and NuoF are missing. This could be due to its low-quality genome, as it consists of 1656 scaffolds with 39.7% of scaffold length in gaps (43). Despite this, we identified its hydrogenase homolog (protein ID 19901 in MycoCosm, with high identity [90%] to that of *C. churrovis*). Therefore, this analysis confirmed that hydrogenosomal Hyd, NuoE, and NuoF are commonly shared among the AF.

**Table 3.** Hydrogenosomal NuoE, NuoF, and Hyd are conserved in AF. The genome of *Pecoramyces sp.* C1A is stored in IMG/MER (genome ID 2510917007) and MycoCosm. The rest 10 AF genomes are from the MycoCosm. The genome of *Piromyces sp.* E2 has 15561 gaps in its genome, accounting for about 40% of scaffold length in gaps. Despite this, the gene encoding the Hyd of *Piromyces sp.* E2 is identified (protein ID 19901). * is for gene ID in IMG/MER.

| Family | Genus | AF Species | Protein ID on MycoCosm |  |  |
| --- | --- | --- | --- | --- | --- |
|  |  |  | NuoE | NuoF | Hyd |
| *Anaeromycetaceae* | *Anaeromyces* | *Anaeromyces robustus* S4 | 267914 | 276695 | 285380 |
| *Caecomycetaceae* | *Caecomyces* | *Caecomyces churrovii* A | 454874 | 557447 | 243125 |
| *Neocallimastigaceae* | *Neocallimastix* | *Neocallimastix constans* G3 | 559368 /<br>674566 | 554324 /<br>671614 | 893170 /<br>902696 |
| *Neocallimastigaceae* | *Neocallimastix* | *Neocallimastix* sp. GF-Ma3-1 | 376615 /<br>705250 | 372829 /<br>547082 | 30729 / 320288 |
| *Neocallimastigaceae* | *Neocallimastix* | *Neocallimastix lanati* | 1083241 /<br>993995 | 1047445 /<br>923302 | 626241 /<br>1341048 |
| *Neocallimastigaceae* | *Neocallimastix* | *Neocallimastix californiae* G1 | 506003 /<br>708584 | 380774 / 51681 | 287975 /<br>377056 |
| *Neocallimastigaceae* | *Neocallimastix* | *Neocallimastix* sp. WI3-B | 810719 /<br>897158 | 273459 /<br>482632 | 483304 /<br>939290 |
| *Neocallimastigaceae* | *Pecoramyces* | *Pecoramyces* sp. F1 | 4429 | 15117 | 16998 |
| *Neocallimastigaceae* | *Pecoramyces* | *Pecoramyces* sp. C1A (*Pecoramyces ruminantium*) | 1184707 | 2511049297* | 2511058566* |
| *Piromycetaceae* | *Piromyces* | *Piromyces* sp. UH3-1 | 529601 | 468346 | 290045 |
| *Piromycetaceae* | *Piromyces* | *Piromyces finnis* | 587149 | 581398 | 585840 |

## DISCUSSION

Hydrogenosomal H_2_ production plays a role in maintaining redox balance in anaerobic eukaryotic cells. The enzymes involved in trichomonad hydrogenosomes H_2_ production are known (9); however, the metabolic pathways for H_2_ production in other anaerobic eukaryotes, such as AF, remain incompletely understood (27, 28), despite its importance in agriculture, biogeochemical cycles in the deep biosphere, and industrial applications. Recent genomic analysis of AF suggested several possibilities for hydrogenosomal H_2_ production (1, 13), but to date, none of them had been experimentally tested due to difficulties with the cultivation of strictly anaerobic fungi and reconstitution of their enzymology. In this study, by combining genomic, metabolic, and proteomic analysis with enzyme assays, we demonstrate that hydrogenosomes of anaerobic fungus *C. churrovis* use a distinct strategy in which electrons from NADH are directly coupled to H_2_ formation through a non-bifurcating NADH-dependent enzyme composed of [FeFe] hydrogenase and NADH dehydrogenase (complex I) subunits NuoE and NuoF.

The identification of a NADH-dependent hydrogenase system in anaerobic fungi provides new insight into the metabolic diversity of mitochondria-derived organelles. Hydrogenosomes have traditionally been viewed as organelles that couple pyruvate oxidation to H_2_ formation through ferredoxin-dependent pathways, as demonstrated in the hydrogenosomes of *T. vaginalis* (9). Our results indicate that AF possess an alternative mechanism in which electrons from NADH are directly linked to H_2_ production through the Hyd–NuoEF system. This pathway provides a route for regenerating NAD^+^ during fermentation and suggests that components related to mitochondrial complex I have been adapted to support anaerobic metabolism in fungi. The widespread occurrence of Hyd and NuoEF homologs in AF genomes indicates that this mechanism may be common across the phylum Neocallimastigomycota.

Our results demonstrate that hydrogenase couples with NuoEF to form H_2_ directly from NADH independently of ferredoxin. The H^+^/H_2_ couple has midpoint redox potential about −414 mV under the standard conditions (25°C, pH 7, 10^5^ Pa), while the midpoint potential of NAD^+^/NADH is about −320 mV at these conditions, making it thermodynamically unfavorable to form H_2_ directly from NADH (44). To make the forward reaction possible, the dissolved H_2_ activity should be maintained low or the ratio of NADH to NAD^+^ should be maintained high to drive the reaction toward H_2_ production. When the dissolved H_2_ activity corresponds to 1-10 Pa, the midpoint potential of H^+^/H_2_ is shifted to around −260 to −300 mV (pH 7 and 25°C) or −279 to −296 mV (pH 7 and 39°C), making the production of H_2_ directly from NADH thermodynamically feasible (45). Several studies reported experimentally verified non-bifurcating NADH-dependent hydrogenase responsible for H_2_ production in bacteria (e.g. *S. wolfei* and *Syntrophus aciditrophicus*) (31, 42). The use of a non-bifurcating NADH-dependent hydrogenase requires a low dissolved H_2_ activity in the aqueous phase. To maintain a low dissolved H_2_ activity, *S. wolfei* and *S. aciditrophicus* live with a H_2_-consuming partner (*i.e.* hydrogenotrophic methanogens) during growth on fatty and alicyclic acids (31). Similar to these syntrophic organisms, AF have a well-known relationship with H_2_-consuming rumen bacteria and methanogens in their native environment (e.g. rumen) (15, 46–48). These H_2_-consuming prokaryotes help keep a low dissolved H_2_ activity by consuming H_2_, changing the metabolic profile of AF (49). Another way to overcome the thermodynamic hurdle is to couple an endergonic reaction with an exergonic reaction. Electron-confurcating hydrogenase generates H_2_ using both reduced ferredoxin and NADH. It drives endergonic redox reaction (synthesis of H_2_ at the expense of NADH) by coupling it to exergonic redox reaction (synthesis of H_2_ at the expense of reduced ferredoxin), effectively conserving the chemical energy (50). Considering the energy benefits conferred by electron confurcation/bifurcation, recent reports speculate that such a mechanism could occur in the hydrogenosomes (1, 10, 13). However, so far there is no experimental evidence of such a mechanism in the hydrogenosomes of anaerobic eukaryotes, and our experiments ruled out this possibility in the hydrogenosomes of anaerobic fungus *C. churrovis*.

Our results indicate that ferredoxin does not contribute to H_2_ production in AF hydrogenosomes. The ferredoxin needs to be first reduced to donate electrons for H_2_ formation. However, we detected no activities of enzymes known to reduce ferredoxin. The recombinant fungal ferredoxin was not reduced with pyruvate by the large organelle fraction samples, by NADH with purified NuoEF, or by H_2_ with the hydrogenase-NuoEF mixture. We observed no activity of PFOR in the large organelle fraction samples, despite proteomic detection of PFOR homolog. The activity of PFOR was observed in *Neocallimastix* sp. L2 and *N. patriciarum* (5, 16), but was undetectable in *N. frontalis* (33). Future work must characterize this enzyme in AF to resolve ongoing debates (5, 16, 33). Ferredoxin in hydrogenosomes and mitochondria supplies electrons for iron-sulfur cluster assembly machinery (51). Unexpectedly, our proteomic analysis of *C. churrovis* did not detect the presence of the predicted hydrogenosomal ferredoxin, yet we expressed recombinant CcFd for assays. Interestingly, ferredoxin was also absent in the proteome of *S. wolfei*, which harbors a functional non-bifurcating NADH-dependent hydrogenase (42, 52).

Heterologous expression of active eukaryotic [FeFe] hydrogenases in *E. coli* hosts has been challenging because of codon bias, oxygen sensitivity, complex maturation process, and improper folding or assembly (53, 54). Despite these challenges, we generated active *C. churrovis* hydrogenase. However, our co-purified Hyd-NuoE-NuoF complex has no H_2_:MV oxidoreductase activity, implying low maturation efficiency in the cell overexpressing multiple proteins at once, including the *S. oneidensis* MR-1 hydrogenase maturases, Hyd, NuoE, and NuoF. Overexpression of NuoE and NuoF might interfere with the maturation process of hydrogenase in the same host cell, because active hydrogenase was successfully generated and purified when co-expressing with hydrogenase maturases in the absence of NuoE and NuoF. By producing Hyd and NuoEF individually, isolating them, and mixing them under *in vitro* conditions, we were able to generate functional enzymes forming H_2_ from NADH. This approach could be useful in studying other oxygen-sensitive proteins from non-model eukaryotes.

Previous reports identified several differences in the β-subunits of bacterial trimeric non-bifurcating hydrogenases and bifurcating hydrogenases, including protein sequence lengths, number of Fe-S clusters, and key amino acids compositions (31, 42, 55, 56). We found that AF NuoF has features that are similar to the β-subunit of non-bifurcating NADH-dependent [FeFe] hydrogenases rather than those of bifurcating hydrogenases (Supplementary Figure 10). In addition, these AF NuoF protein sequences are shorter than the β subunit of bifurcating hydrogenases, and they do not have the extra two Fe-S clusters that are proposed to be interacting with ferredoxin in the β-subunit of bifurcating hydrogenases (31, 55, 56). Similarly, the monomeric [FeFe] hydrogenase of *Nyctotherus ovalis* has features similar to that of AF NuoF and β-subunit of non-bifurcating NADH-dependent hydrogenase in bacteria (31). It was suggested to be a non-bifurcating NADH-dependent hydrogenase (11, 12); however, the biochemical evidence is still missing. Recently, it was proposed that hydrogenosomes of trichomonads (e.g. *T. vaginalis*) (1, 10) and the mitochondrion-related organelles in some marine anaerobic ciliates may use electron-bifurcating hydrogenase to form H_2_ (57). The hydrogenosomal NuoF of trichomonads shares features with that of non-bifurcating enzymes (similar protein lengths, Fe-S cluster numbers and key amino acids) (Supplementary Figure 10), suggesting that such a bifurcating mechanism may not be used by their hydrogenosomes. Future experimental work will help clarify this issue. Detailed structural characterization of these enzymes can be helpful to elucidate the functional mechanism of non-bifurcating NADH-dependent enzymes.

Understanding the mechanism of H_2_ production in AF allows genetic engineering to redirect carbon flux toward desired fermentation products by modulating H_2_ production. Our work elucidates this mechanism of H_2_ production in AF, identifying new targets for metabolic engineering to optimize AF fermentation. AF produce acetate, ethanol, lactate, and formate, in addition to H_2_. H_2_ production influences carbon flux and product distribution by competing with other NADH-consuming pathways in AF. For example, in co-culture with a hydrogenotrophic methanogen, the methanogen scavenged H_2_ from AF, thereby increasing acetate, succinate, and ethanol yields, but decreasing lactate yield from AF (49). This experimentally verified hydrogenosomal non-bifurcating NADH-dependent hydrogenase pathway could be integrated into the previously published genome-scale metabolic model of AF (13), enhancing its predictive power. Therefore, this discovery not only expands our understanding of eukaryotic hydrogenosomal metabolism, but also has broad implications for bioengineering of H_2_-producing systems.

## MATERIALS AND METHODS

### Organisms

*C. churrovis* was isolated by the O’Malley lab (35). *P. brevis* GA33, *P. ruminicola* 23, and *S. oneidensis* MR-1 (ATCC 700550) were obtained from the ATCC. *C. pasteurianum* W5 was obtained from DSMZ. *T. vaginalis* T1 strain was isolated by J.H.Tai (58). *E. coli* BL21(DE3)Δ*iscR* was a gift from Dr. Thomas Happe (59). *E. coli* NEB5α was from NEB Company.

### Media and growth

*C. churrovis* was grown in anaerobic medium MB with glucose or MC minus with glucose under oxygen-free CO_2_ and with serum bottles with butyl rubber stoppers at 39°C. MB, MC minus medium, and MC medium were made according to literature (49) with modifications (see details in Supplementary Methods). MC medium was used for revival of cryopreserved *C. churrovis* and for culturing of *P. brevis* GA33 and *P. ruminicola* 23. *S. oneidensis* MR-1 was growing in the ATCC medium 18 (Tryptic Soy Broth) at 30°C. *C. pasteurianum* W5 was cultured on a glucose medium at 37°C (60). *E. coli* strains were cultured on Luria-Bertani (LB) medium for cloning work and Terrific Broth (TB) medium for protein expression experiments at 37°C. *T. vaginalis* T1 was growing in the 10 mL of TYM medium supplemented with 10% inactivated horse serum at 37°C (61).

### H_2_ production from *C. churrovis*

*C. churrovis* was regularly transferred to fresh MC minus medium with 5 g/L glucose after 3 to 4 days growth. To grow cells for H_2_ production measurement, 2 mL of the 3-days-old culture was inoculated to 40 mL of fresh medium in a 70-mL serum bottle. One mL of medium was taken for high-performance liquid chromatography (HPLC) analysis before the inoculation. Another 1 mL of culture was withdrawn immediately after the inoculation. Supernatants from the 1 mL medium and 1 mL culture were saved for HPLC analysis. Gas pressure in the bottle was released to 0 psi after inoculation. No H_2_ was detected in the headspace before the incubation at 39°C. Gas pressure and H_2_ production measurement were done on a daily basis using the method described below. After 7 days of growth, culture supernatant was collected for HPLC analysis.

*C. churrovis* cultures were shaken manually right before the measurement of gas pressure and H_2_ production. Gas pressure in the bottle headspace was measured by a pressure meter. The headspace gas pressure was released to 0 psi while the released gas volume was measured and collected by a syringe. H_2_ amount in the headspace (30 mL) was measured by injecting 100 µL gas sample to gas chromatograph (Thermo TRACE 1300) for analysis using the H_2_ detection method described in the literature (49) and calibrated with standard H_2_ gas. The released H_2_ gas amount was calculated by multiplying H_2_ concentration measured by the gas chromatography and the volume of gas collected in the syringe. The accumulated H_2_ amount is the sum of H_2_ in the headspace and in the syringe. Four cultures were included as independent replicates.

### HPLC analysis

Glucose and several major fermentation products in the supernatant of *C. churrovis* culture were measured by HPLC by modifying the method described in the literature (49). Specifically, glucose, formate, and ethanol signal were monitored by refractive index detector. Acetate, lactate, and succinate were monitored using a variable wavelength detector set to 210 nm.

### Preparation of large organelle fraction (LOF) of *C. churrovis*

After 3 days growth on MC minus medium with 5 g/L glucose, 3 mL culture was inoculated to 40 mL same medium. After 2 days growth, 2 bottles of such culture were pooled and harvested under N_2_ atmosphere. The following steps were done under N_2_ atmosphere. Cells were harvested at 3000 × *g*, 4°C for 5 min, and washed with 16 mL anaerobic buffer (20 mM Tris-HCl [pH 7.5], 2 mM dithiothreitol [DTT], and 250 mM sucrose). Cell pellet was resuspended in 8 mL same buffer, sonicated for 5 min with 3 s on and 6 s rest at 30% amplitude (Branson 250 sonicator). The lysate was spun at 1000 × *g*, 4°C for 10 min to get pellet and supernatant. The pellet was resuspended in a 10 mL same buffer and centrifuged. Supernatants from the two centrifugations were pooled into a new centrifuge tube. One and half mL of 50% (w/v) OptiPrep in anaerobic buffer (250 mM sucrose, 1 mM EDTA, 10 mm Tris-HCl [pH 7.4]) was gently injected to the tube bottom via a long needle. After centrifugation at 18,000 × *g*, 4°C for 30 min, supernatant was collected as the cytosol fraction. The thin organelle layer on top of OptiPrep was mixed with the OptiPrep and 8 mL same buffer, and centrifuged at 12,000 × *g*, 4°C for 10 min. The pellet containing large organelle fraction (LOF) was gently resuspended in 2 mL buffer and aliquoted. These fractions (cytosol, LOF) were kept in air-tight screw tubes. An aliquot of these organelles were mixed with Triton X-100 (final concentration 2% [v/v]) by pipetting and used for enzyme assays. Remaining untreated organelles were freshly frozen in liquid nitrogen and stored at −80°C until use. LOF aliquot was quickly thawed at 39°C water bath, kept on ice, and ready for use in protein concentration measurement and enzyme assays. At least three biological replicates of LOF samples were prepared.

### Isolation of *C. churrovis* hydrogenosome fractions

After 3 days growth on MB medium with 5 g/L glucose, 2 mL culture was inoculated to 40 mL same medium. Cells were harvested after 3 to 4 days growth at 3000 × *g* for 5 min at room temperature, washed with 50 mM potassium phosphate buffer (KP_i_, pH 6.4), weighted, and frozen at −20°C until use. Cell pellets harvested from three independently grown cultures were used as biological replicates. About 5 g of wet cells were used to isolate hydrogenosomes from *C. churrovis*. For each 1 g cells, 0.4 mg chitinase (Sigma-Aldrich #C6137) and 5 mg β-glucanase (Sigma-Aldrich # G4423) were dissolved in 5 mL of 50 mM KP_i_ (pH 6.4). They were mixed with cell pellet and incubated at 39°C water bath for 5 min followed by 1 h incubation at 39°C in a shaking incubator to weaken cell wall. Cell resuspension was centrifuged at 2500 × *g*, 4°C for 5 min to get pellet. Each gram of cell pellet was resuspended in 2.5 mL buffer (20 mM KP_i_ [pH 7.4] and 250 mM sucrose). Each of the 1.25 mL cell resuspension was mixed with 0.8 mL zirconia/silica beads (BioSpec #11079105Z) in a 2-mL screw tube and kept on ice before lysis (3 cycles of 10 s on and 15 s rest on ice) by a beadbeater (BioSpec Mini-beadbeater 16), followed by 5 min maximum-speed vortex on a vortex mixer (Fisher Scientific #02215365). Cell lysate and beads mixture were pooled and centrifuged at 2500 × *g*, 4°C for 10 min. The supernatant was centrifuged at 8000 × *g*, 4°C for 20 min to get organelles pellet, which was resuspended in 1.5 mL buffer (20 mM KP_i_ [pH 7.4] and 250 mM sucrose) and loaded to the top of OptiPrep gradients (18% - 50%, w/v) for ultracentrifugation at 25,000 RPM for 4 h at 4°C in a swing-out rotor (Beckman Coulter #SW28) using a Beckman Coulter ultracentrifuge Optima XE-90. The OptiPrep gradients contained 3 mL of 18% OptiPrep on the top, 3 mL 20%, 22%, 24%, 26%, 28%, 30%, 32%, 34%, 36%, and 1.5 mL 50% OptiPrep at the bottom. These gradients were made by diluting 60% (w/v) OptiPrep into buffer (60 mM Tris-HCl [pH 7.4], 250 mM sucrose) to get 50% OptiPrep, which was further diluted in buffer (10 mM Tris-HCl [pH 7.4], 1 mM EDTA, and 250 mM sucrose) to get various lower concentrations. Various fractions were formed after ultracentrifugation and collected by long needles. Each fraction was diluted with 5 volumes of the buffer (20 mM Tris-HCl [pH 7.5], 250 mM sucrose, 0.5 mM KCl) and centrifuged at 16,000 × *g*, 4°C for 15 min. The pellets were resuspended in 0.25 mL of the same buffer, aliquoted, and stored at −80°C until use.

### Preparation of *P. brevis* GA33 cytoplasmic fraction and crude hydrogenase of *C. pasteurianum* W5

*P. brevis* GA33 cytoplasmic fraction was made according to the literature (60) with modifications (see Supplementary Methods for detailed method). Cells were lysed by sonication under N_2_ atmosphere. Unbroken cells and cell debris were removed by centrifugation at 18,000 × *g*, 4°C for 10 min and cell membrane was removed by ultracentrifugation at 20,800 × *g*, 4°C for 1 h. Crude hydrogenase of *C. pasteurianum* W5 was prepared according to the literature (40).

### Cloning and protein purification

Overall, genes were amplified from genome DNA, boiled cell lysate, or synthesized DNA, using the Phusion PCR Master Mix (NEB #M0531L). Plasmid backbones were prepared by PCR, digested with DpnI, and gel-purified. These DNA fragments were assembled using a Gibson Assembly Kit (NEB #E2621L). The correct sequences of plasmids were verified by sequencing at Plasmidsaurus Inc. They were transformed into *E. coli* BL21(DE3)Δ*iscR* for protein overexpression. Plasmid pACYCDuet1-HydGX-HydEF was co-transformed with pET32a-Hyd-Strep or pET32a-Strep-Hyd-NuoE-NuoF into *E. coli* BL21(DE3)Δ*iscR*. After inducing protein overexpression and harvesting the cells, TvFd and CcFd were purified under aerobic conditions, while recombinant OGOR, Hyd-Strep, NuoEF-Strep, Strep-Hyd-NuoE-NuoF were purified under anaerobic conditions (under N_2_ atmosphere or in the anaerobic chamber), by modifying methods in the literature (62, 63). These Strep-tag II fusion proteins were purified via affinity chromatography. Protein concentration was measured with the Pierce Bradford Plus protein assay reagent (Thermo Scientific #23238) using bovine serum albumin as standard protein to create a calibration curve. Details about plasmids, *E. coli* strains, cloning, and protein purification methods are listed in Table 4, Table 5, and Supplementary methods.

**Table 4.** Expression plasmids used in this work.

| Plasmid | Protein | Tags | Induction |
| --- | --- | --- | --- |
| pASKIBA7-Strep-TvFd | Matured TvFd of *T. vaginalis* | N-term Strep-Tag II | anhydrotetracycline |
| pASKIBA7-Strep-OGOR | OGOR of *P. ruminicola* 23 | N-term Strep-Tag II | anhydrotetracycline |
| pET32a-CcFd-Strep | Matured CcFd of *C. churrovis* | C-term Strep-Tag II | IPTG |
| pET32a-Hyd-Strep | Matured Hyd of *C. churrovis* | C-term Strep-Tag II | IPTG |
| pET32a-NuoEF-Strep | Matured NuoE and NuoF of *C. churrovis* | C-term Strep-Tag II of NuoF | IPTG |
| pET32a-Strep-Hyd-NuoE-NuoF | Matured Hyd, NuoE, and NuoF of *C. churrovis* | N-term Strep-Tag II of Hyd | IPTG |
| pACYCDuet1-HydGX-HydEF | HydE, HydF, and HydG of *S. oneidensis* MR-1 | Not applicable | IPTG |

**Table 5.**
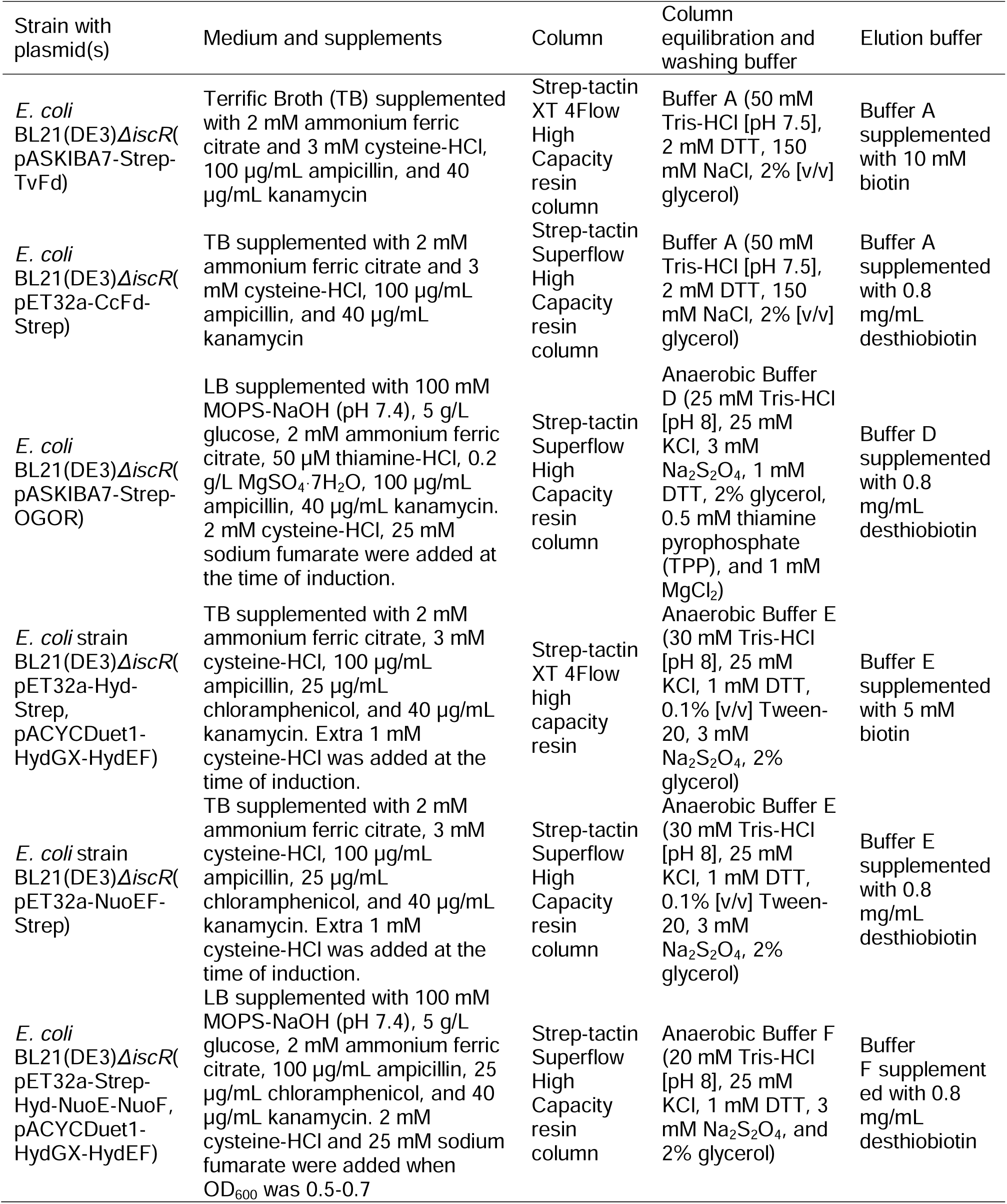
Protein overexpression and purification conditions in this study.

### Enzyme assays

We performed enzyme assays with the Triton X-100 treated large organelle fraction (LOF) samples and purified proteins under N_2_ atmosphere in anaerobic cuvettes with anaerobically prepared reagents unless specified. We used semi-micro absorption cuvettes (Hellma USA #114-10-40) capped with grey butyl rubber stopper (DWK Life Sciences Wheaton # W224100405) for most enzyme assays unless specified. When H_2_ was used as a substrate, we used a home-made anaerobic cuvette capped with blue butyl rubber stopper. These cuvettes are made of a cuvette (Hellma USA #100-10-40) and glass tubing with two glass balls on its side. Protein samples were added to the side balls before mixing with other reagents in the cuvette. Enzyme assays were performed at room temperature unless specified. Absorbance was monitored throughout the reaction at 340 nm for NAD(P)H, 578 nm for reduced methyl viologen (MV), 420 nm for CcFd, and 460 nm for TvFd. Molar extinction coefficient for NADH and NADPH is 6220/M/cm. Molar extinction coefficient for MV is 11,100/M/cm. Based on absorbance change before and after reduction by crude hydrogenase with *C. pasteurianum* W5 with H_2_, we calculated the molar extinction coefficient of our purified CcFd and TvFd to be 2895/M/cm and 2525/M/cm, respectively. Assay components were added to the assay in the order as listed in the assay components. The maximum reaction rates after initiation of the reaction were used to calculate its specific activity. One unit (1 U) is defined as the amount of enzyme that catalyzes conversion of 1 µmol of a substrate per minute under specific conditions.

The enzyme assay for malic enzyme (NAD^+^ or NADP^+^) activity (0.5 mL) contained 100 mM Tris-HCl (pH 7.8), 10 mM MgCl_2_, 15 mM KCl, 0.5 mM NAD^+^ or NADP^+^, LOF sample (about 4-7 µg), and 10 mM malic acid. Malic acid was added last to initiate the reaction.

The 1-mL enzyme assay for pyruvate:MV oxidoreductase activity contained 50 mM Tris-HCl (pH 7.6), 10 mM MgCl_2_, 4 mM DTT, 0.2 mM CoA, 0.4 mM TPP, 2 mM MV, LOF sample (about 58-82 µg), and 10 mM pyruvate. The enzyme assay for pyruvate:CcFd oxidoreductase activity was the same as that for pyruvate:MV oxidoreductase except that we used 37 µM CcFd rather than 2 mM MV. At the end of this enzyme assay, we spiked about 58 µg cytoplasmic fraction protein of *P. brevis* GA33 to the assay to show that this enzyme assay worked for measuring pyruvate:MV oxidoreductase and pyruvate:CcFd oxidoreductase activity.

The enzyme assays for 2-oxoglutarate:MV oxidoreductase, 2-oxoglutarate:TvFd oxidoreductase, 2-oxoglutarate:CcFd oxidoreductase activity (0.5 mL) contained 50 mM Tris-HCl (pH 7.6), 10 mM MgCl_2_, 4 mM DTT, 0.2 mM CoA, 0.4 mM TPP, 2 mM MV or 50.7 µM TvFd or 51.6 µM CcFd, 6.5 µg purified OGOR, and 10 mM 2-oxoglutarate.

The enzyme assay for NADH:MV oxidoreductase activity (0.5 mL) contained 25 mM Na-glycine (pH 8.5), 5 mM MV, 10 mM KCl, 0.4 mM NADH, and LOF sample (about 6-10 µg) or purified proteins (about 0.35-0.57 µg). LOF sample or purified protein was added last to initiate the reaction. The enzyme assay for NADPH:MV oxidoreductase activity (0.5 mL) contained 25 mM Na-glycine (pH 8.5), 5 mM MV, 10 mM KCl, 2 mM NADPH, and purified protein (350-570 ng). The enzyme assay for NADH:CcFd oxidoreductase and NADH:TvFd oxidoreductase activity (0.5 mL) contained 25 mM Na-glycine (pH 8.5), 60 µM CcFd or TvFd, 10 mM KCl, 0.4 mM NADH, and about 350 ng purified protein.

The enzyme assay for H_2_:MV oxidoreductase activity (1 mL) contained 25 mM Na-glycine (pH 8.5), 10 mM DTT, 1 mM MV, and LOF sample (about 58-82 µg) or purified proteins (1.1-2.6 µg) or crude hydrogenase (2.5 µg) under H_2_ atmosphere. These components were added to the cuvette under N_2_ atmosphere, while the sample(s) was placed into a side ball. The assay was then bubbled with H_2_ gas for 5-10 min. Protein sample was mixed with the assay to initiate the reaction.

The enzyme assay for H_2_:NAD^+^ oxidoreductase or H_2_:NADP^+^ oxidoreductase activity (1 mL) contained 25 mM Na-glycine (pH 8.5), 10 mM DTT, 1 mM NAD^+^ or 1 mM NADP^+^, 10 mM KCl, 5 µM FMN (flavin mononucleotide), and LOF sample (about 58-82 µg) or purified enzymes (5 µg) under H_2_ atmosphere. The assay was performed in the same way as that for measuring H_2_:MV oxidoreductase activity but absorbance was monitored at 340 nm.

To measure if H_2_:NAD^+^ oxidoreductase activity in LOF sample is dependent on CcFd, we added CcFd after the NAD^+^ reduction reaction reached a plateau. Briefly, the 1 mL enzyme assay contained CcFd (37 µM) and other components were the same as that for H_2_:NAD^+^ oxidoreductase assay. Absorbance at 340 and 420 nm was monitored. Protein sample and CcFd were separately placed at two side balls before mixing with the assay. The reaction was initiated by mixing with the LOF sample. When the rate of NAD^+^ reduction reached a plateau, we mixed CcFd with the assy. To measure if H_2_:NAD^+^ oxidoreductase activity in purified proteins is dependent of CcFd or TvFd, we performed the same assay as that for LOF sample but used 5 µg proteins (mixture of 2.5 µg Hyd-Strep and 2.5 µg NuoEF), 30 µM CcFd and TvFd.

The enzyme assay for H_2_:CcFd oxidoreductase and H_2_:TvFd oxidoreductase activity (1 mL) contained 25 mM Na-glycine (pH 8.5), 10 mM DTT, protein sample (about 6 µg Hyd-Strep), and 30 µM CcFd or TvFd under H_2_ atmosphere. Protein sample and CcFd or TvFd were placed in the two side balls before mixing with the assay. When the absorbance slope rate reached a plateau after adding protein samples, CcFd or TvFd was then mixed with the assay. The enzyme assays to measure these activities in crude hydrogenase of *C. pasteurianum* W5 (Cp5-Hyd) were similar to that for Hyd-Strep, but 2.5 µg Cp5-Hyd, and 60 µM TvFd or 51.6 µM CcFd were used.

### H_2_ production assay

#### H_2_ production from NADH or NADPH

The 2-mL enzyme assay was done in a 7.5 mL serum bottle capped with blue butyl rubber stopper under N_2_ atmosphere at 39°C. A mini-stir bar was placed inside the serum bottle. The enzyme reaction was done at 39°C with constant magnetic stirring at 300 RPM. The enzyme assay contained 100 mM KP_i_ (pH 7.4), 10 mM DTT, 5 µM FMN, enzyme samples (about 4.5 µg Hyd-Strep and 5.7 µg NuoEF-Strep), and 2 mM NADH or NADPH. The assay was incubated at 39°C for 15 min before adding enzyme samples and NADH or NADPH. Enzyme samples were mixed for 5 min at room temperature under N_2_. The reaction was started by addition of enzyme samples and NADH or NADPH. A hundred µl of gas sample from the headspace were taken immediately after that and also every 8 to 10 min. H_2_ in the gas sample was analyzed by gas chromatography. Anaerobic water was added in a control assay without NADH or NADPH.

#### H_2_ production from reduced ferredoxin and NADH

The 2-mL assay contained 100 mM KP_i_ (pH 7.4), 10 mM DTT, 5 µM FMN, a ferredoxin-reducing system, enzyme samples (about 4.5 µg Hyd-Strep and 5.7 µg NuoEF-Strep), and 2 mM NADH. The reaction was done in a 7.5 mL serum bottle capped with a butyl rubber stopper under N_2_ atmosphere at 39°C. Ferredoxin-reducing system contained 0.5 mM MgCl_2_, 0.2 mM CoA, 0.4 mM TPP, various amount of ferredoxin (25.8 µM CcFd, or 15.2 µM TvFd, or 5.7 µM Cp5Fd (ferredoxin purified from *Clostridium pasteurianum* W5) (34), and 19.2 µg recombinant OGOR, and 10 mM 2-oxoglutarate. The assay components were mixed in the order listed above. The assay was incubated at 39°C for 30 min to reduce ferredoxin by OGOR with 2-oxoglutarate before adding enzyme samples and NADH. Enzyme samples (Hyd-Strep and NuoEF-Strep) were mixed for 5 min at room temperature and then added to the assay. Gas samples (100 µL) from the headspace were taken at every 8 to 10 min interval. About 30-35 min later after adding enzyme samples to the reaction, we added 2 mM NADH to the assay. Anaerobic water was added to a control assay without ferredoxin. We performed control assays to verify that the ferredoxin-reducing system can reduce ferredoxin to donate electrons for H_2_ production. The control assay contained 100 mM KP_i_ (pH 7.4), 10 mM DTT, 5 µM FMN, ferredoxin-reducing system with 25.8 µM CcFd, or 15.2 µM TvFd, or 2.8 µM Cp5Fd. The reaction was initiated by adding crude hydrogenase of *C. pasteurianum* W5 (about 29-200 µg).

### Enzyme assays with various organelle fractions

We measured malic enzyme (NADP^+^) activity with the 8 different fractions (#1-8) to identify which fraction(s) contained hydrogenosomes. These enzyme assays (aerobic) were done in a 96-well plate (flat clear bottom). The 100 µL malic enzyme (NADP^+^) assay contained 100 mM Tris-HCl (pH 7.8), 10 mM MgCl_2_, 15 mM KCl, 0.5 mM NADP^+^, 0.2% (v/v) Triton X-100, 2.4 µg organelle fraction resuspended in 40 µL 2% (v/v) Triton X-100 in 20 mM Tris-HCl (pH 7.5) and 250 mM sucrose, and 10 mM potassium malate. Absorbance at 340 nm was monitored by a plate reader (Agilent Biotek Synergy H1). The assays (200 µL) for comparing activity of malic enzyme (NAD^+^) and malic enzyme malic enzyme (NADP^+^) contained 100 mM Tris-HCl (pH 7.8), 10 mM MgCl_2_, 15 mM KCl, 0.5 mM NAD^+^ or NADP^+^, 2.4 µg organelle fraction resuspended in 40 µL 2% (v/v) Triton X-100 in 20 mM Tris-HCl (pH 7.5) and 250 mM sucrose, and 10 mM potassium malate.

### SDS-PAGE, native PAGE, western blot

Protein samples were mixed with 4x Laemmli Sample Buffer (Bio-Rad #1610747), heated at 95°C for 5 min, and loaded to a precast gel for SDS-PAGE analysis. Gel was visualized by staining with Coomassie G-250. To run a native PAGE, we mixed protein samples with 4x NativePage Loading Buffer (Invitrogen #BN2003) and loaded them to a 4-16% precast gel (Invitrogen #BN1002BOX). Electrophoresis was performed according to manual instructions.

We confirmed the correct expression of Strep-tag II proteins via western blot. Briefly, the purified proteins were denatured by boiling, electrophoresed, and transferred to a 0.22 µm PVDF membrane (Bio-Rad # 1704156) using a defined protocol in a Bio-Rad Trans-Blot Turbo Transfer System. The membrane was washed with 1xTBST (20 mM Tris base, 150 mM NaCl, 0.1% [v/v] Tween 20) for 5 min, incubated in 1xTBST containing 5% (w/v) nonfat milk for 1 h, and incubated in 1xTBST containing 5% (w/v) nonfat milk and 1 x Precision Protein StrepTactin-HRP Conjugate (Bio-Rad #1610381) for 1 h. The membrane was washed with 1xTBST for 5 min for 5 times. Strep-tagged proteins were visualized by Pierce ECL western blot substrates (Thermo Scientific #32209) and the Bio-Rad ChemiDoc imaging system.

### NanoPOTS analysis of isolated hydrogenosome fractions

We performed enzyme assays with the fractions (#1-8) collected after OptiPrep density gradient centrifugation. The fractions with high malic enzyme activity (NADP^+^) (#4, 5, 6) were pooled and processed as one sample. In total, three replicates of independently prepared fractions were used for nanoPOTS as described in the literature (37) (see details in Supplementary Methods).

### Global proteomic analysis of cell lysate of *C. churrovis*

Four replicates of cells were harvested after 3 days of growth in MC minus with 5 g/L glucose. Cell pellets were flash frozen in liquid nitrogen and stored at −80°C. Sample protein extraction from the cell pellets was based on the previous protocol (64). Peptide digests were analyzed using a data-dependent acquisition (DDA) approach on liquid chromatography-tandem mass spectrometry (LC-MS/MS) platform (see Supplementary Methods). LC-MS/MS data was searched against the *C. churrovis* proteome database using the MS-GF+ search tool (65). Peptide intensities were subsequently extracted using MASIC software (66).

### LC-MS/MS analysis of purified proteins and proteins in gel band

We used untargeted proteomics to identify proteins in the purified samples (Hyd-Strep, NuoEF-Strep). The two batches of independently purified Hyd-Strep were pooled together for analysis according to literature(67). NuoEF-Strep samples were analyzed in the same way.

We used LC-MS/MS to identify proteins in the native PAGE gel. The gel band was cut into pieces and destained by washing in 25 mM ABC for 5 min, 50% (v/v) acetonitrile and 25 mM ABC for 5 min, and 100% acetonitrile. We repeated this step and then washed them in 100 mM ABC for 15 min, 100% acetonitrile for 10 min, 100 mM ABC for 15 min, 100% acetonitrile for 10 min. The dried gel pieces were incubated with 100 mM DTT and 50 mM ABC for 90 min at 56°C. Excess liquid was removed and gel pieces were shrank in 100% acetonitrile, and alkylated in 55 mM iodoacetamide for 20 min in the dark. The excess liquid was removed and gel pieces were washed with 50 mM ABC for 15 min twice, shrank with 100% acetonitrile, and digested with trypsin in 150 µL of 50 mM ABC at 37°C overnight. The digestion reaction was quenched by 6.5 µL formic acid. The peptides were extracted, analyzed by LC-MS/MS in DDA mode, and identified from LC-MS/MS data as described in literature (34).

### Bioinformatic analysis

We identified *C. churrovis* hydrogenase (Hyd), NuoE, NuoF, CcFd by searching for protein annotations in MycoCosm. Homologs of these proteins in other AF were identified by the BLAST tool on MycoCosm using protein sequences of Hyd, NuoE, and NuoF of *C. churrovis* as input. The genome of *Pecoramyces sp.* C1A in IMG/M (IMG genome ID 2510917007) was searched to identify genes encoding Hyd, NuoE, and NuoF and compared to that in MycoCosm. We used the BLASTP algorithm on NCBI website to analyze identities between protein sequences (https://blast.ncbi.nlm.nih.gov/Blast.cgi). NCBI CD-search (https://www.ncbi.nlm.nih.gov/Structure/cdd/wrpsb.cgi) was used to predict cofactor preference of the malic enzyme of *C. churrovis* (protein ID 462551 on MycoCosm). Protein localization was predicted by DeepLoc2 (68). We predicted structures of *C. churrovis* matured Hyd, NuoE, and NuoF with cofactors NAD^+^ and FAD (flavin-adenine dinucleotide) in the AlphaFold Serve (https://alphafoldserver.com/) using default setting. The predicted structure was visualized by ChimeraX (69). Protein structures of Hyd1ABC of *S. wolfei* (IMG/M Locus Tag ID Swol_1017, Swol_1018, Swol_1019) were predicted in the same way.

## Statistics

A one-sided t-test was used to determine if mean yield of fermentation products and mean values of enzymatic activity was greater than 0. This test was also used to determine if the rate of H_2_ production from enzyme assays was greater than 0, and if H_2_ production rates were different in the assays compared to that with only NADH.

## Data availability

The nanoPOTS mass spectrometry data and processed results have been uploaded to MassIVE with the accession MSV000100685. The mass spectrometry proteomics data for heterologously generated proteins and proteins in gel bands have been deposited to the ProteomeXchange Consortium via the PRIDE (70) partner repository with the dataset identifier PXD070885 (Project accession).

## ACKNOWLEDGEMENTS

We are grateful to Dr. Timothy Hackmann at UC Davis and Dr. Thomas Happe at Ruhr University Bochum for sharing research materials. We would like to thank Nguyen Thu Duong, Paulína Pristašová, Tien Le, Michal Havelka, and Jitka Kučerová at the University of Charles in Prague for their technical support in enzyme assays and antiserum generation. We thank Dr. Gabriela Grigorean of the UC Davis Proteomics Core for performing LC-MS analysis of purified proteins and proteins in gel band. The authors acknowledge the use of the Biological Nanostructures Laboratory within the California NanoSystems Institute, supported by the University of California, Santa Barbara and the University of California, Office of the President. This research was partially supported by the DOE Office of Science, Office of Biological and Environmental Research (BER), grant no. DE-SC0022142. This research was further supported by the NSF ExFAB BioFoundry and supported by The National Science Foundation under Award No. DBI-2400327. The research at Charles University (JT, IH) was supported by the Ministry of Education, Youth and Sport of the Czech Republic (Inter-Action LUAUS23052).

## Author Contributions

Conceptualization: B.Z., M.A.O; Methodology: B.Z., I.H., J.T., Y.G., S.M.W., J.M.F., N.M., M.B.; Data curation and formal analysis: B.Z., I.H., J.T., Y.G., S.M.W., J.M.F., N.M., M.B.; Visualization: B.Z., J.M.F; Writing – original draft: B.Z., Y.G., S.M.W., J.M.F., N.M., M.B.; Writing – review & editing: B.Z., I.H., J.T., Y.G., S.M.W., J.M.F., N.M., M.B., S.E.B., M.A.O.; Funding acquisition: M.A.O., J.T., S.E.B.; Project administration: B.Z., M.A.O., S.E.B.

## SUPPLEMENTAL MATERIAL

Supplementary information: Supplementary Methods, Supplementary Table 1-4 and Figure 1-10. Supplementary dataset 1-5.

## Notes

### Competing Interest Statement

The authors have declared no competing interest.

