## Supplementary Methods, Supplementary Table 1-4 and Figure 1-10 for "The anaerobic fungus *Caecomyces churrovis* produces H_2_ via a non-bifurcating NADH-dependent enzyme complex"

**SUPPLEMENTARY METHODS, TABLES, AND FIGURES**

- **Supplementary Methods**
- **Supplementary Table 1-4**
- **Supplementary Figure 1-10**
- **References**

**Supplementary Methods**

**Media**

Per liter, the MB medium contained 100 mL of stock salt solution, 5 mL hemin stock solution, 10 mL ATCC trace mineral supplement (Fisher #50-238-3325), 0.5 mL of 0.1% (w/v) resazurin, 4 g sodium carbonate, 1 g cysteine-HCl, 1 mL of 100 g/L MOPS [3-(N-morpholino)propanesulfonic acid] , 1 mL ATCC vitamin (Fisher Scientific #50-189-789FP), and 12.5 mL of 40% (w/v) glucose solution. The stock salt solution was made of 6.8 g/L KH_2_PO_4_, 6 g/L KCl, 6 g/L NaCl, 5 g/L MgSO_4_·7H_2_O, 2 g/L CaCl_2_·2H_2_O, and 5.4 g/L NH_4_Cl. To make 1 L hemin stock solution, we dissolved 0.1 g hemin in 10 mL ethanol and adjusted total volume to 1 L with deionized distilled water containing 2 g NaOH. To make the medium, we mixed the stock salt solution, hemin stock solution, ATCC trace mineral supplement, resazurin, and about 1.15 L deionized distilled water in a 2.8 L flask, and heated it using microwave to ensure that it was boiled for at least 5 min to remove oxygen. The medium volume was measured and the final volume was adjusted to 1 L with anaerobic water. The boiled medium was bubbled with CO_2_. Sodium carbonate was added while it was cooled to room temperature. After about 15 min, cysteine-HCl was added. The medium was aliquoted to serum bottles under CO_2_ atmosphere and autoclaved at 121°C for 20 min. Anaerobic sterile MOPS, ATCC vitamin solution, and glucose solution were added after autoclaving.

The MC minus medium was made in the similar way as the MB medium. Per liter, the MC minus medium contained 0.25 g yeast extract (Gibco #212750), 0.5 g Bacto Casitone (Gibco #225930), 150 mL of 3 g/L K_2_HPO_4_, 150 mL salt stock solution containing 3 g/L KH_2_PO_4_, 6 g/L (NH_4_)_2_SO_4_, 6 g/L NaCl, and 0.6 g/L MgSO_4_·7H_2_O, 75 mL centrifugation-clarified rumen fluid, 0.2 mL 0.1% (w/v) resazurin, 6 g sodium bicarbonate, 1 g cysteine-HCl and 12.5 mL of 40% (w/v) glucose solution. Anaerobic sterile glucose solution was added after autoclaving at 5 g/L concentration.

MC medium was used for revival of cryopreserved *C. churrovis* and for culturing of *P. brevis* GA33 and *P. ruminicola* 23. Compared to the MC minus medium, the MC medium has higher amounts of yeast extract, Casitone and rumen fluid (2.5 g yeast extract, 10 g Bacto Casitone, and 150 mL centrifugation-clarified rumen fluid) per liter. Glucose was added after autoclaving.

**Preparation of *P. brevis* GA33 cytoplasmic fraction and crude hydrogenase of *C. pasteurianum* W5**

*P. brevis* GA33 was growing in 40 mL MC medium with 5 g/L glucose under CO_2_ atmosphere at 39°C until stationary phase. Half mL of the stationary-phase culture was used as seed to inoculate 40 mL medium. Four bottles of culture were inoculated and incubated at 39°C for 16 h. The following steps were performed under N_2_ atmosphere. These cultures were pooled and harvested by centrifugation at 16,000 × *g* for 5 min at 4°C. Cells were washed twice in the anaerobic buffer containing 50 mM Tris-HCl (pH 7.6), 20 mM MgSO_4_, 4 mM DTT, and 10% (v/v) glycerol. The cell pellet was resuspended in 16 mL of the same buffer supplemented with 0.2 mM PMSF and 25 U/mL Benzonase Nuclease (Sigma-Aldrich #E1014), and sonicated for a total 10 min with 10 s on, 20 s off, and 30% amplitude under N_2_ atmosphere (Branson 250). The lysate was centrifuged at 18,000 × *g* for 10 min at 4°C and the supernatant was further centrifuged at 20,800 × *g* for 1 h at 4°C. The final supernatant (the cytoplasmic fraction) was aliquoted to 2-mL air-tight screw tubes (SARSTEDT #72.694.107), frozen in liquid nitrogen, and then stored at -80°C until use.

Crude hydrogenase of *C. pasteurianum* W5 was made based on the published procedure (1). Briefly, 20 mL culture at stationary phase were inoculated into 450 mL of a glucose medium in a 1-L Pyrex bottle capped with bromobutyl rubber stopper (DWK Life Sciences #292062803) and incubated at 37°C for 46 h(2). Two bottles of 450 mL medium were inoculated. Gas generated during growth was released into the atmosphere. The cells were harvested by centrifugation at 12,000 × *g* for 3 min at 4°C and washed twice with 16 mL anaerobic buffer (50 mM Tris-HCl, pH 8). The cell pellet (about 4.3 g wet weight) was resuspended in 8.6 mL same buffer. The cell resuspension (about 13.5 mL) was supplemented with 0.5 mg/mL lysozyme and 12.5 U/mL Benzonase Nuclease, and incubated at 37°C for 50 min with occasional mixing. The lysate was centrifuged at 7000 × *g* for 20 min at 4°C and the supernatant was transferred to a serum bottle under N_2_ atmosphere. The supernatant was then bubbled with H_2_ for 5 min, capped with butyl stopper, and incubated at 55°C for 15 min under H_2_ atmosphere. It was then incubated on ice for 15 min and transferred to an air-tight centrifuge tube for centrifugation at 18,000 × *g* for 30 min at 4°C. This supernatant was aliquoted into 2-mL air-tight tubes under N_2_ and stored at -80°C until use. Its hydrogenase activity was detected by enzyme assays.

**Plasmids construction**

Overall, genes were amplified from genome DNA, boiled cell lysate, or synthesized DNA, using the Phusion PCR Master Mix (NEB #M0531L). Plasmid backbones were prepared by PCR, digested with DpnI, and gel-purified. These DNA fragments were assembled using a Gibson Assembly Kit (NEB #E2621L). The correct sequences of plasmids were verified by sequencing at Plasmidsaurus Inc.

Gene encoding the matured ferredoxin 1 (TvFd) of *Trichomonas vaginalis* (ferredoxin 1 without leader peptide MLSQVCRF, TVAG_003900 in the TrichDB database) was cloned into plasmid pASK-IBA7 at the site downstream of Strep-tag II, generating plasmid pASKIBA7-Strep-TvFd. Genome DNA of 10 mL overnight culture of *T. vaginalis* was isolated using a DNA extraction kit (Geneaid #GB300) and used to amplify TvFd DNA fragment. Genes encoding 2-oxoglutarate:ferredoxin oxidoreductase (OGOR) of *Prevotella ruminicola* 23 (PRU_2267, PRU_2268, Locus Tag in IMG/MER) were inserted into pASK-IBA7 at the site downstream of Strep-tag II, generating plasmid pASKIBA7-Strep-OGOR. Cells in 100 µL *P. ruminicola* 23 culture in stationary phase were pelleted, resuspended in 100 µL sterile water, and boiled for 10 min. Cell resuspension was centrifuged and the supernatant was used to amplify sequences containing PRU_2267, PRU_2268, and the segment of nucleotides in between.

To clone the genes encoding possible ferredoxin (CcFd), NADH dehydrogenase subunit E and F (NuoE, NuoF), and hydrogenase (Hyd) of *C. churrovis*, we identified their protein IDs in MycoCosm (549362 for CcFd, 454874 for NuoE, 557447 for NuoF, 243125 for Hyd). We predicted their possible leader sequences using the DeepLoc2 (3). These genes encoding matured proteins (without leader sequences) and the DNA sequences for peptide containing Strep-tag II (SAWSHPQFEK) were codon optimized and synthesized by IDT (Integrated DNA Technologies, Inc.). The sequence between NuoE and NuoF is TTTGTTTAACTTTAAGAAGGAGATATACAT. We amplified these DNA fragments and inserted them into pET32a to replace the sequence between the start codon (ATG) and the stop codon (TAA) within the sequence downstream of the ribosome binding site but upstream of the T7 terminator. This generated plasmids pET32a-Hyd-Strep, pET32a-NuoEF-Strep, and pET32a-CcFd-Strep. The Strep-tag II was located at the C-terminal of the protein(s) of interest. Similarly, the plasmid pET32a-Strep-Hyd-NuoE-NuoF was constructed so that the DNA sequence between the start codon (ATG) and the stop codon (TAA) of pET32a was replaced by the sequence (from 5’ to 3’) containing the DNA fragments for Hyd with Strep-tag II (ASWSHPQFEK) located at its N-terminal, CCTCTAGAAATAATTTTGTTTAACTTTAAGAAGGAGATATACAT, NuoE, TTTGTTTAACTTTAAGAAGGAGATATACAT, and then NuoF. No Strep-tag II was added to NuoE or NuoF.

We cloned the four genes encoding HydG (SO3923, Locus Tag on IMG/M), HydX (SO3924), HydE (SO3925), HydF (HM357716, Accession ID on NCBI) of *Shewanella oneidensis* MR-1 into plasmid pACYCDuet-1 based on literature (4). The hydrogenase maturases are HydE, HydF, and HydG, but HydX is encoded by the HydGXEF operon with no indentified functions. The genome DNA was extracted via a QIAGEN kit (Qiagen #51804) and used as template for amplifying these genes. The genes encoding HydG and HydX were assembled into the first multiple cloning site of pACYCDuet-1, generating pACYCDuet1-HydGX. These genes replaced the DNA sequence between the start codon (ATG) and stop codon (TAA) including the His-tag region in the pACYCDuet-1. The genes encoding HydE and HydF were assembled with the backbone of pACYCDuet1-HydGX to generate pACYCDuet1-HydGX-HydEF. They replaced the sequence between ATG and TAA including the S-tag sequence of the second multiple cloning site of pACYCDuet1-HydGX.

**Protein overexpression and purification**

The plasmids mentioned above were used to transform *E. coli* BL21(DE3)*ΔiscR* for protein overexpression. The pASKIBA7 and pET32a derived plasmids confer cells with ampicillin resistance. pACYCDuet-1 derived plasmids enable cells to be resistant to chloramphenicol. *E. coli* BL21(DE3)*ΔiscR* is resistant to kanamycin. pACYCDuet1-HydGX-HydEF was co-transformed with pET32a-Hyd-Strep or pET32a-Strep-Hyd-NuoE-NuoF into *E. coli* BL21(DE3)*ΔiscR*. After harvesting the cells, TvFd and CcFd were purified under aerobic conditions, while recombinant OGOR, Hyd-Strep, NuoEF-Strep, Strep-Hyd-NuoE-NuoF were purified under anaerobic conditions (under N_2_ atmosphere or in the anaerobic chamber), by modifying methods in the literature (5, 6). Protein concentration was measured with the Pierce Bradford Plus protein assay reagent (Thermo Scientific #23238) using bovine serum albumin as standard protein to create a calibration curve.

*E. coli* strain BL21(DE3)*ΔiscR*(pASKIBA7-Strep-TvFd) was inoculated to the Terrific Broth medium (TB) supplemented with 2 mM ammonium ferric citrate, 3 mM cysteine-HCl, 100 µg/mL ampicillin, and 40 µg/mL kanamycin and grown at 37°C. When optical density at 600 nm (OD_600_) of the culture reached to 0.6-0.9, 200 ng/mL anhydrotetracycline was added to induce expression of TvFd. After induction at 37°C for about 7 h, cells were harvested by centrifugation at 12,000 × *g* for 5 min at 4°C, washed with Buffer A (50 mM Tris-HCl [pH 7.5], 2 mM DTT, 150 mM NaCl, 2% [v/v] glycerol), and resuspended in Buffer A for lysis by sonication (total 40 min with cycles of 3 s on and 6 s rest at 30% amplitude). Cell lysate was spun at 18,000 × *g* for 30 min at 4°C and the supernatant was spun at 50,000 × *g* for 20 min at 4°C. Supernatant was loaded to a Strep-tactin XT 4Flow High Capacity resin column for protein purification, following the instructions (IBA LifeSciences 25030010). The purified TvFd was mixed with Buffer C (50 mM Tris-HCl [pH 7.5], 150 mM NaCl, 2% [v/v] glycerol) and concentrated with a 10-kDa cut-off filter. The concentrated TvFd was aliquoted, frozen in liquid nitrogen, and stored at -80°C until use. We generated CcFd using the same method as that for TvFd with modifications. Briefly, 0.5 mM IPTG was used to induce protein expression in *E. coli* BL21(DE3)*ΔiscR*(pET32a-CcFd-Strep). The buffer for cells lysis contained Buffer A supplemented with 0.2 mM PMSF and 4 U/mL Benzonase Nuclease. A Strep-tactin Superflow High Capacity resin column was used to purify CcFd following manual instructions (IBA LifeSciences 21208010).

*E. coli* BL21(DE3)*ΔiscR*(pASKIBA7-Strep-OGOR) was inoculated to LB medium supplemented with 100 mM MOPS-NaOH (pH 7.4), 5 g/L glucose, 2 mM ammonium ferric citrate, 50 µM thiamine-HCl, 0.2 g/L MgSO_4_·7H_2_O, 100 µg/mL ampicillin, and 40 µg/mL kanamycin. Cells were growing at 37°C at 200 RPM until the OD_600_ reached 0.5-0.7. The culture was brought into an anaerobic chamber (5% H_2_, 20% CO_2_, 75% N_2_) and stirred with a stir bar. We added 2 mM cysteine-HCl, 25 mM sodium fumarate, and 200 ng/mL anhydrotetracycline to the culture to induce protein production. After growing overnight at room temperature in the anaerobic chamber, cells were harvested by centrifugation, washed with anaerobic Buffer D (25 mM Tris-HCl [pH 8], 25 mM KCl, 3 mM Na_2_S_2_O_4_, 1 mM DTT, 2% glycerol, 0.5 mM thiamine pyrophosphate (TPP), and 1 mM MgCl_2_), and resuspended in Buffer D. About 40 mL cell resuspension was lysed by sonication under N_2_ atmosphere (15 min on, with cycles of 10 s on and 20 s off at 30% amplitude). The cell lysate was spun at 18,000 × *g* for 15 min at 4°C and the supernatant was passed through 0.22 µm filter. It was loaded to Strep-Tactin Superflow High Capacity resin column. Protein purification was performed in the anaerobic chamber following the manual instructions. Buffer D was used to equilibrate and wash the column. The elution buffer was Buffer D supplemented with 0.8 mg/mL desthiobiotin. Purified proteins were aliquoted into 2-mL screw tubes, quickly frozen in liquid nitrogen, and kept at -80°C until use.

*E. coli* strain BL21(DE3)*ΔiscR*(pET32a-Hyd-Strep, pACYCDuet1-HydGX-HydEF) was cultured in TB supplemented with 2 mM ammonium ferric citrate, 3 mM cysteine-HCl, and appropriate antibiotics, at 37°C at 250 RPM. When OD_600_ reached 0.6-0.9, 0.5 mM IPTG and 1 mM cysteine-HCl were added. The culture was bubbled with N_2_ for 2 h at room temperature and transferred into a Pyrex bottle capped with butyl rubber stopper for growth overnight at room temperature. Cells were harvested, washed with a freshly-made Buffer E (30 mM Tris-HCl [pH 8], 25 mM KCl, 1 mM DTT, 0.1% [v/v] Tween-20, 3 mM Na_2_S_2_O_4_, 2% glycerol), and resuspended in the Buffer E supplemented with 0.8 mM PMSF and 4 U/mL Benonase Nuclease for sonication. Cell lysate was spun at 50,000 × *g* for 20 min at 4°C. The supernatant was loaded to Strep-tactin XT 4Flow high capacity resin column for purification following the manual instructions. Buffer E was used to equilibrate and wash the column. Buffer E supplemented with 5 mM biotin was for eluting Hyd-Strep. Eluted fractions were collected into 2-mL screw tubes, fresh frozen in liquid nitrogen, and kept at -80°C until use.

The overexpression and purification of NuoEF-Strep were the same as that for Hyd-Strep, except that NuoEF-Strep was purified via the Strep-Tactin Superflow high capacity resin column. The elution buffer was the Buffer E supplemented with 0.8 mg/mL desthiobiotin.

*E. coli* BL21(DE3)*ΔiscR*(pET32a-Strep-Hyd-NuoE-NuoF, pACYCDuet1-HydGX-HydEF) was used to generate Strep-Hyd, NuoE, and NuoF. Cells were grown in the LB supplemented with 100 mM MOPS-NaOH (pH 7.4), 5 g/L glucose, 2 mM ammonium ferric citrate, and appropriate antibiotics. After OD_600_ reached 0.5-0.7, 2 mM cysteine-HCl and 25 mM sodium fumarate were added, and the culture was bubbled with N_2_ for 30 min before addition of 0.5 mM IPTG. Culture was placed in anaerobic chamber. A sterilized stir bar was placed in the culture for stirring at room temperature. Cells were harvested, washed with Buffer F (20 mM Tris-HCl [pH 8], 25 mM KCl, 1 mM DTT, 3 mM Na_2_S_2_O_4_, and 2% glycerol), and resuspended in the same buffer supplemented with 0.2 mM PMSF for sonication. Cell lysate was spun at 18,000 × *g* for 15 min at 4°C. Supernatant was filtered through a 0.22 µm filter and loaded to a Strep-Tactin Superflow High Capacity resin column for protein purification. After loading lysate to the equilibrated column, the column was washed with 10 column volumes (CV) of Buffer F. Proteins were eluted with Buffer F supplemented with 0.8 mg/mL desthiobiotin). Seven fractions of 0.5 CV of elutes were collected and stored at -80°C until use.

**NanoPOTS analysis of isolated hydrogenosome fractions**

Samples for LC-MS analysis were removed from -80°C in PCR tubes and placed on ice prior to centrifugation at 16,000 × *g* for 15 min at 4°C. The pellet was resuspended in 10 μL of extraction buffer (25 mM ammonium bicarbonate [ABC], 0.5 x PBS, 0.1% n-dodecyl-β-D-maltoside [DDM], 2 mM DTT) and incuated at 70°C for 1 h at 1000 RPM. These samples were centrifuged at 1000 × *g* for 30 s and 1 μL of 60 mM chloroacetamide was added before incubation in the dark for 30 minutes. Ten microliters of solution containing 16 ng/µL trypsin and 8 ng/µL Lys-C in 50 mM ABC were added, followed by overnight digestion at 37°C at 1,500 RPM in a ThermoMixer. Quenching was accomplished with addition of 1 μL of 5% formic acid and samples were centrifuged for 15 min at 6000 × *g* at 4°C. Aliquots of 20 µL peptide solution were placed in low-volume polypropylene LC-MS vials with MicroSolv caps and stored at -20°C. Prior to LC-MS analysis, these samples were diluted 1:1 with 0.1% formic acid in LC-MS grade water.

We employed a Waters Acquity M Class UPLC for nanoflow chromatographic separation. For trapping, a custom packed SPE column (150 μm i.d., 4 cm, 5 μm particle size, 300 Å pore size C18 material, Phenomenex) was used and the analytical LC column (75 μm i.d., 35 cm long, 1.7 μm particle size, 190 Å pore size C18 material, Waters) was made in-house with a self-pack picofrit (cat. no. PF360-50-10-N-5, New Objective, Littleton, MA). The analytical column was heated to 50 °C using a 15-cm AgileSleeve column heater (Analytical Sales and services, Inc., Flanders, NJ). Briefly, trapping was performed on the SPE column at 4 µL/min for 5 min. After washing the peptides, samples were eluted at 0.2 µL /min and separated using a 60 min gradient with Buffer B (0.1% formic acid in acetonitrile).

For mass spectrometry analysis, a Bruker timsTOF-SCP with a Captive Spray source was coupled to our in-house nanoPOTS autosampler. The timsTOF-SCP was operated in high-sensitivity, DDA-PASEF mode with a duty cycle of 1.56 s consisting of 8 PASEF MS/MS scans from 100 to 1700 m/z with an ion mobility range (1/K0) from 0.6 to 1.6 Vs/cm^2^. Capillary voltage was set to 1500 V. The TIMS ramp and accumulation time were set to 166 ms while the collision energy was ramped linearly as a function of mobility from 59ev at 1/K_0_ = 1.6 Vs/cm^2^ to 20 eV at 1/K_0_ = 0.6 Vs/cm^2^. Precursors with charge state from 0 to 5 were selected with a target value of 10,000 and intensity threshold of 500. Isolation width was set to 2 m/z at or below 700 m/z and 3 m/z at or above 800 m/z, with a linear interpolation between the two points. Isolated precursors were excluded from analysis for 0.4 min after isolation.

All Bruker “.d“ files were processed by FragPipe (version 20.0) and searched against the *C. churrovis* protein sequence database (14,772 protein entries) acquired from the Joint Genome Institute. These databases included target sequences as well as decoy sequences and common protein contaminants. MSfragger version 3.8, IonQuant version 1.9.8, and Philosopher version 5.0.0 were applied to the search. Search settings included a precursor mass tolerance of +/- 20 ppm, fragment mass tolerance of +/- 20 ppm, deisotoping, trypsin enzyme specificity, carbamidomethylation as a fixed modification, and several variable modifications (oxidation of methionine, N-terminal acetylation, and pyro-glutamate). Protein and peptide identifications were filtered to a false discovery rate of less than 1% within FragPipe. IonQuant match-between-runs (MBR) was set to “TRUE” and an MBR false discovery rate of 1% at ion level was used to reduce false positive matches. Peptide abundances were rolled up to protein abundances using a top N strategy and median normalization across runs was performed within FragPipe. For comparing abundances of different proteins, intensity Based Absolute Quantification (iBAQ) values were calculated that normalize for a protein’s size based on the number of theoretical tryptic peptides. These values were calculated by summing all peptide intensities for a given protein and dividing it by the number of theoretically observable tryptic peptides between 6 and 50 amino acids in length (7).

**Global proteomic analysis of cell lysate of *C. churrovis***

Four replicates of samples were collected in this experiment. For each replicate, 0.5 mL of 2-days age culture seed (*C. churrovis*) was inoculated to 40-mL medium MC minus with 5 g/L glucose. Gas pressure was monitored and released after each day. After 3 days of growth, the culture was harvested via centrifugation at 3000 × *g* for 5 min at 4°C. The cells were washed twice with anaerobic buffer containing 50 mM Tris-HCl (pH 7) and 2 mM DTT. Cell pellets were flash frozen in liquid nitrogen and stored at -80°C. Sample protein extraction from the cell pellets was based on the previous protocol (8). Peptide digests were analyzed using a data-dependent acquisition (DDA) approach on liquid chromatography-tandem mass spectrometry (LC-MS/MS) platform, including a Thermo Vanquish Neo UHPLC coupled with a Thermo Q Exactive HF-X mass spectrometer. A total of 500 ng of peptides per sample were first loaded onto a primary trap column (Jupiter 5 μm, 150 μm i.d.), operated at 5.5 μL/min flow rate using 0.1% formic acid in water as the loading solution. Peptides were then separated on a reverse-phase Waters AQ C18 column (1.7 μm, 30 cm, 75 μm i.d.), maintained at 45°C with a controlled flow rate of 0.2 μL/min. The mobile phases used for separation consisted of (A) 0.1% formic acid in water and (B) 0.1% formic acid in acetonitrile, and a gradient was applied over 182 minutes: 0 min, 1%B; 4 min, 1%B; 14 min, 8%B; 108 min, 21%B; 133 min, 28%B; 145 min, 37%B; 151 min, 75%B; 155 min, 95%B; 161 min, 95%B; 163 min, 50%B; 164 min, 1%B; 165 min, 95%B; 166 min, 1%B; 182 min, 1%B. Following separation, peptides were introduced into the mass spectrometer via electrospray ionization. Full MS survey scans were performed across a mass range of 300 to 1800 m/z, with a resolution of 60,000, an AGC target of 3e6, and a maximum injection time (IT) of 20 ms, to select precursor ions based on intensity. The Top 12 precursor ions were then selected for fragmentation using a 0.7 m/z isolation window with dynamic exclusion enabled for 45 seconds. Fragmented was carried out using high-energy collisional dissociation (HCD) at a normalized collision energy (NCE) of 30, and the fragment ions were recorded in MS/MS scans with a mass range of 200 to 1200 m/z, a resolution of 45,000, an AGC target of 1e5, and a maximum IT of 100 ms.

LC-MS/MS data was searched against the *C. churrovis* proteome database using the MS-GF+ search tool (9). The search parameters included dynamic oxidation of methionine residues and static carbamidomethylation of cysteine residues. A partially tryptic search was performed with a parent ion tolerance set to 20 ppm. Systematic mass measurement bias was calibrated and corrected using the mzRefinery tool (10). Search results were filtered to retain only peptide-spectrum matches (PSMs) with mass errors between -5 ppm and 5 ppm (i.e., -5< DelM_PPM < 5) and 1% false discovery rates (FDRs) (i.e., Q-value < 0.01). Proteins identified with only single-peptide hits were excluded from the final dataset. Peptide intensities were subsequently extracted using MASIC software (11).

**Supplementary Table 1**. Major fermentation products formed by *C. churrovis* grown in the medium MC minus with 5 g/L glucose. Four cultures were included as biological replicates. Glucose was consumed during growth. SEM, Standard Error of the Mean.

|  | µmol (in 40 mL culture) | |
| --- | --- | --- |
|  | Mean | SEM |
| H_2_ | 106.1 | 2.9 |
| Glucose | 767.1 | 21.9 |
| Formate | 715.7 | 10.9 |
| Ethanol | 146.4 | 3.8 |
| Acetate | 359.7 | 6.4 |
| Lactate | 605.0 | 1.9 |
| Succinate | 78.2 | 1.5 |

**Supplementary Table 2.** Malic enzyme (NAD^+^ or NADP^+^) detected in the isolated hydrogenosomes fractions. Their enzyme activities are shown as Mean ± SEM mU/mg protein.

|  | malic enzyme (NAD^+^)  (mU/mg protein) | malic enzyme (NADP^+^)  (mU/mg protein) |
| --- | --- | --- |
| #4 | 352 ± 113 | 100 ± 31 |
| #5 | 277 ± 52 | 86 ± 27 |
| #6 | 287 ± 91 | 78 ± 22 |

**Supplementary Table 3.** Peptides of Hyd, NuoE, NuoF, PFOR, and ME of *C. churrovis* were detected by global proteomic analysis*.* Count represents the number of biological replicates where the peptide was detected among four replicates. In total, 48 peptides for Hyd, 17 peptides for NuoE, 26 peptides for NuoF, 9 peptides for PFOR, and 108 peptides for ME were detected.

| **Peptide** | **Count** | **Peptide** | **Count** | |
| --- | --- | --- | --- | --- |
| **Hyd (protein ID 243125)** | | **ME (protein ID 462551)** | | |
| KNLNPEDIIHVSVMPCTAK | 2 | FTTDEKDR | | 4 |
| MCPGGCINGGGQPK | 2 | PSEVYNKK | | 2 |
| EEHPNDCM*TCESNGNCEFQDLIYR | 2 | ETIVDCIVAEGATR | | 4 |
| PMDRPMNFTK | 2 | LGAGSSGVGVCETIVDCIVAEGATR | | 1 |
| QDMSLHQK | 3 | AYAQFYMFDHK | | 4 |
| AIQDMSLHQK | 3 | APSEVYNKK | | 4 |
| ANGFYIPTLCYHPR | 3 | ASGSPFDPVEYK | | 4 |
| CVECGQCSQVCPVGAITER | 4 | ETFQVCAR | | 2 |
| FPMFTSCCPGWINMVEK | 4 | FAPSEVYNKK | | 3 |
| GKNGEVVPDIDYVLTTR | 4 | IYLNHLQNR | | 4 |
| GSWKPLTACTTEVWEGMEIETDTPTVR | 3 | PGIGLGLVSCR | | 4 |
| INPSELQNDKYDSPLGIGSSAGNLFGVTGGVMEAAVR | 4 | PLSNPTSR | | 4 |
| KINPSELQNDKYDSPLGIGSSAGNLFGVTGGVMEAAVR | 4 | TGGNLIFASGSPFDPVEYK | | 4 |
| KTNHSITEPCYSPFDNSTFSIAR | 3 | TTDEKDR | | 4 |
| KTTVLDAAK | 4 | YMFDHK | | 4 |
| MVAGLR | 4 | AITLASVLATMR | | 2 |
| MVNISINGR | 4 | GTFADIKK | | 4 |
| NGEVVPDIDYVLTTR | 4 | YLNHLQNR | | 4 |
| NLNPEDIIHVSVMPCTAK | 2 | AVIQFEDFM*M*PNALDLLLK | | 3 |
| RPEFTR | 3 | AVIQFEDFMMPNALDLLLK | | 4 |
| SPQQMMGAVIK | 3 | DQICMFNDDIQSTGAITLASVLATMR | | 4 |
| TNHSITEPCYSPFDNSTF | 4 | DYQGGKTPAEMLK | | 1 |
| TNHSITEPCYSPFDNSTFSIAR | 4 | ETFLCLGAGSSGVGVCETIVDCIVAEGATR | | 4 |
| TTVLDAAK | 3 | FIYLNHLQNR | | 4 |
| TYFAQK | 3 | FIYLNHLQNRNETLYYK | | 3 |
| TYFAQKK | 3 | GRDDLLPSQSVFM*R | | 3 |
| VIVATTAPAVR | 3 | GRDDLLPSQSVFMR | | 3 |
| FTSCCPGWINMVEK | 1 | GSAFTTDEK | | 4 |
| IDAQHPVR | 4 | GSAFTTDEKDR | | 4 |
| AAVVSGGANIQK | 4 | HNM*WQPEYPHIVVK | | 4 |
| ACHHFQNINILGFINR | 4 | HNMWQPEYPHIVVK | | 4 |
| EEHPNDCMTCESNGNCEFQDLIYR | 4 | IFTQTR | | 4 |
| KVIVATTAPAVR | 4 | IKEPLEK | | 3 |
| LCLVDVK | 4 | KETFLCLGAGSSGVGVCETIVDCIVAEGATR | | 1 |
| LNEGGKFPMFTSCCPGWINMVEK | 4 | KQVIVTKK | | 4 |
| LPIAGNCR | 1 | KTGLDIINDPK | | 4 |
| LSHENPEITQIYK | 2 | LNKGSAFTTDEK | | 3 |
| MYTMDEEAK | 4 | LTLYVCGGGINPR | | 4 |
| NECIEVLR | 3 | M*ILENFVELAPIIYTPVVGEACQK | | 4 |
| SALAEEYNADPTFDFTGK | 4 | MILENFVELAPIIYTPVVGEACQK | | 4 |
| SSLAMMR | 3 | QVIVTKK | | 4 |
| TAQVITGVENPIPLGELK | 3 | TGLDIINDPK | | 3 |
| TAQVITGVENPIPLGELKAIRG | 4 | TIQTNQCNNSYSFPGIGLGLVSCR | | 4 |
| VGTPM*DRPMNFTK | 2 | TPAEM*LK | | 3 |
| VGTPMDRPM*NFTK | 4 | TPAEMLK | | 2 |
| VGTPMDRPMNFTK | 2 | YKDQICM*FNDDIQSTGAITLASVLATMR | | 4 |
| YQIDAQHPVR | 4 | YKDQICMFNDDIQSTGAITLASVLATM*R | | 3 |
| GTPMDRPMNFTK | 3 | YKDQICMFNDDIQSTGAITLASVLATMR | | 4 |
|  |  | CLGAGSSGVGVCETIVDCIVAEGATR | | 3 |
| **NuoE (protein ID 454874)** | | GLGDLGAGGMQIPIGK | | 2 |
| FTPENLK | 2 | IFASGSPFDPVEYK | | 3 |
| FTPENLKR | 4 | PSQSVFMR | | 4 |
| KYPDNFKR | 3 | TLYVCGGGINPR | | 3 |
| TGLTALTSEPTGPR | 4 | VPPRPQSLEAQYQR | | 4 |
| VCGSDEIFNTIK | 3 | FNDDIQSTGAITLASVLATMR | | 4 |
| VCGSDEIFNTIKR | 4 | ILENFVELAPIIYTPVVGEACQK | | 4 |
| YLLQVCGTTPCK | 2 | SAVAANWPYDDVDVIVVTDGSR | | 4 |
| GATIPLLDIAQR | 4 | YFSTADR | | 3 |
| GKYLLQVCGTTPCK | 1 | FVELAPIIYTPVVGEACQK | | 3 |
| KGVVPK | 1 | NSYSFPGIGLGLVSCR | | 3 |
| VDAKPSNTAPFK | 4 | QCNNSYSFPGIGLGLVSCR | | 2 |
| VYEVATFYTMFHR | 2 | SYSFPGIGLGLVSCR | | 4 |
| LSSTPIVAR | 2 | PRPQSLEAQYQR | | 4 |
| DAKPSNTAPFK | 4 | PEYPHIVVK | | 4 |
| TM*FHREPR | 2 | SLEAQYQR | | 2 |
| ELGINIGETTK | 2 | ARGGTFADIKK | | 4 |
| VYEVATFYTMFHREPR | 2 | DDLLPSQSVFM*R | | 2 |
|  |  | DDLLPSQSVFMR | | 3 |
|  |  | EEAYAQFYM*FDHK | | 4 |
| **NuoF (protein ID 557447)** | | EEAYAQFYMFDHK | | 4 |
| STFAGSKPLTDEDRIWTNLYGR | 4 | EEAYAQFYMFDHKG | | 3 |
| GGWDNLLAVIPGGASCPLIPR | 3 | EEAYAQFYMFDHKGLLGKG | | 4 |
| GQDWIINEMK | 3 | GGTFADIK | | 4 |
| LFCISGCVNNPCTVEESMGISMR | 4 | GGTFADIKK | | 4 |
| LIEGILIAGR | 4 | GLVPPRPQ | | 3 |
| NACGSGWDMDIYIQR | 4 | GLVPPRPQSL | | 3 |
| WFSSFGR | 4 | GLVPPRPQSLEAQYQR | | 4 |
| WSFLNKPLDGR | 4 | GM*YFSTADR | | 4 |
| WSFLNKPLDGRPR | 2 | GMYFSTADR | | 4 |
| YQAAHPK | 3 | GMYFSTADRGQM | | 3 |
| AVQAHTSYIYIR | 3 | GQM*SAVAANWPYDDVDVIVVTDGSR | | 3 |
| EGTGWMAEMMER | 4 | GQMSAVAANWPYDDVDVIVVTDGSR | | 4 |
| GAGAYVCGDETALIESIEGK | 4 | ILGLGDLGAGGM*QIPIGK | | 3 |
| GEFVNEAETLQK | 4 | ILGLGDLGAGGMQIPIGK | | 4 |
| GGAGFATGLK | 4 | KDYQGGKTPAEM*LK | | 2 |
| GRGGAGFATGLK | 4 | KDYQGGKTPAEMLK | | 4 |
| HESCGQCTPCR | 1 | M*NIATQFPPK | | 4 |
| IWTNLYGR | 4 | MNIATQFPPK | | 4 |
| LEVGNM*TIEELNQLER | 1 | NETLYYK | | 4 |
| LEVGNMTIEELNQLER | 4 | SECTAEEAVEFTGGNLIFASGSPFDPVEYK | | 4 |
| LTYNIEGR | 3 | STRVPFETFQVCAR | | 4 |
| PLADTACMDYDSMSDCHSSLGTGAMLVLDK | 3 | VPFETFQVCAR | | 4 |
| SFRPYCEER | 4 | AITDRFPK | | 3 |
| TICALGEAAALPVR | 3 | FFAPSEVYNK | | 1 |
| VKPPFPADFGVFGCPTIVNNVETISATPTIMR | 4 | FFAPSEVYNKK | | 4 |
| YLVINCDEGEPNTCK | 4 | NLDKIKEPLEK | | 4 |
|  |  | PFETFQVCAR | | 3 |
| **PFOR (protein ID 530611)** |  | PPRPQSLEAQYQR | | 4 |
| FAVLGFGDSSYER | 2 | AQFYMFDHK | | 3 |
| FTVLGLGSTSK | 4 | SFPGIGLGLVSCR | | 4 |
| LLNSLIDIENR | 3 | VCGGGINPR | | 4 |
| SGEYNVHAHEASK | 3 | YAQFYMFDHK | | 2 |
| WIQDILK | 4 | HAFVDEWVSAITDRFPK | | 2 |
| YSIISIEDATK | 2 | MILENFVELAPIIYTPVVGEAC | | 2 |
| AGIPEVVIHAADDVSLDDLKK | 2 | ASVLATMR | | 2 |
| VINIIYGTDLGTTK | 1 | LIFASGSPFDPVEYK | | 2 |
| YAVFGLGDSTR | 2 | AVAANWPYDDVDVIVVTDGSR | | 1 |
|  |  | GEEFHAFVDEWVSAITDRFPK | | 2 |

**Supplementary Table 4.** Specific activity for H_2_ production by the mixture of Hyd-Strep and NuoEF-Strep.

| Reaction | Activity(nmol/min/mg protein)  [Mean (SEM)] | *P*-value to test if activity is different than 0 | *P*-value to test if activity is different than that with NADH |
| --- | --- | --- | --- |
| without NADH | 0 | / | 0.0070 |
| with NADH | 98 (1) | 0.0035 | / |
| with only reduced Fd (CcFd or TvFd or Cp5Fd) | 0 | / | / |
| after adding NADH to the assay without reduced Fd | 79 (1) | 0.0039 | 0.0680 |
| after adding NADH to the assay with reduced CcFd | 75 (2) | 0.0085 | 0.0847 |
| after adding NADH to the assay with reduced TvFd | 86 (11) | 0.0446 | 0.3168 |
| after adding NADH to system with reduced Cp5Fd | 105 (6) | 0.0189 | 0.2925 |
| with NADPH | 0 | / | / |

**Supplementary Figure 1**. Enzyme activities were measured with the large organelle fraction (LOF) of *C. churrovis*. NAD^+^, NADP^+^, MV (methyl viologen), or pyruvate was added to the assay before measuring absorbance for (A-H). The assay was bubbled with H_2_ for 5 min before measuring absorbance for (C-E). (A) Malic enzyme (NAD^+^) activity of LOF. (B) Malic enzyme (NADP^+^) activity of LOF. (C) Addition of LOF resulted in reduction of NAD^+^ with H_2_, showing H_2_:NAD^+^ oxidoreductase activity. (D) Addition of LOF resulted in reduction of NAD^+^ with H_2_ but addition of CcFd didn’t result in higher rate of NAD^+^ reduction. (E) MV was reduced by LOF with H_2_, showing H_2_:MV oxidoreductase activity. (F) NADH:MV oxidoreductase activity of LOF. (G) Addition of LOF and MV to the assay didn’t result in reduction of MV with pyruvate. Addition of a positive control for pyruvate:MV oxidoreductase (cytoplasmic fraction of *P. brevis* GA33, GA33CE) resulted in reduction of MV, showing that the assay works for detecting pyruvate:MV oxidoreductase activity. (H) No CcFd was reduced by LOF with pyruvate. (I) Addition of a positive control (GA33CE) for pyruvate:CcFd oxidoreductase to the reaction in (H) lead to reduction of CcFd, showing that the assay works for detecting pyruvate:CcFd oxidoreductase activity. One representative figure for each assay is shown. Three independently prepared samples (LOF) were tested.

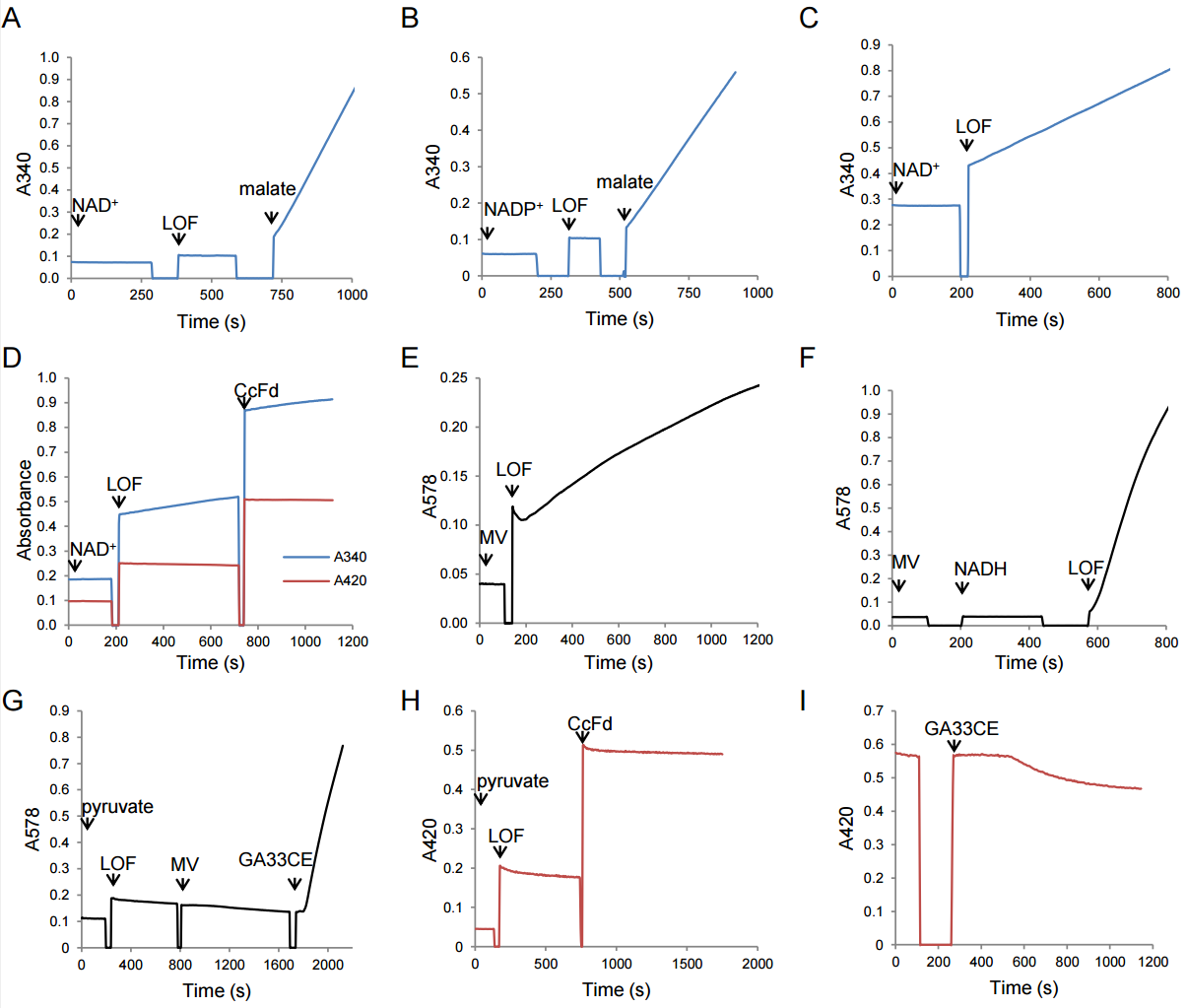

**Supplementary Figure 2**. Western blot confirmed the expression of β subunit of SCS (succinyl-CoA synthetase) (βSCS) in the isolated fractions #4, 5, 6 that were pooled for NanoPOTs proteomic analysis. Primary antibody was generated in rat by injecting it with a heterologously generated SCS β subunit of *Neocallimastix lanati*. Secondary antibody was goat anti-rat IgG H&L (HRP) (Abcam #ab97057). Signal was visualized by a Pierce Kit ECL (Thermo Scientific #32209). β subunit of SCS of *Neocallimastix lanati* shares 96.6% identity with that of *C. churrovis*.

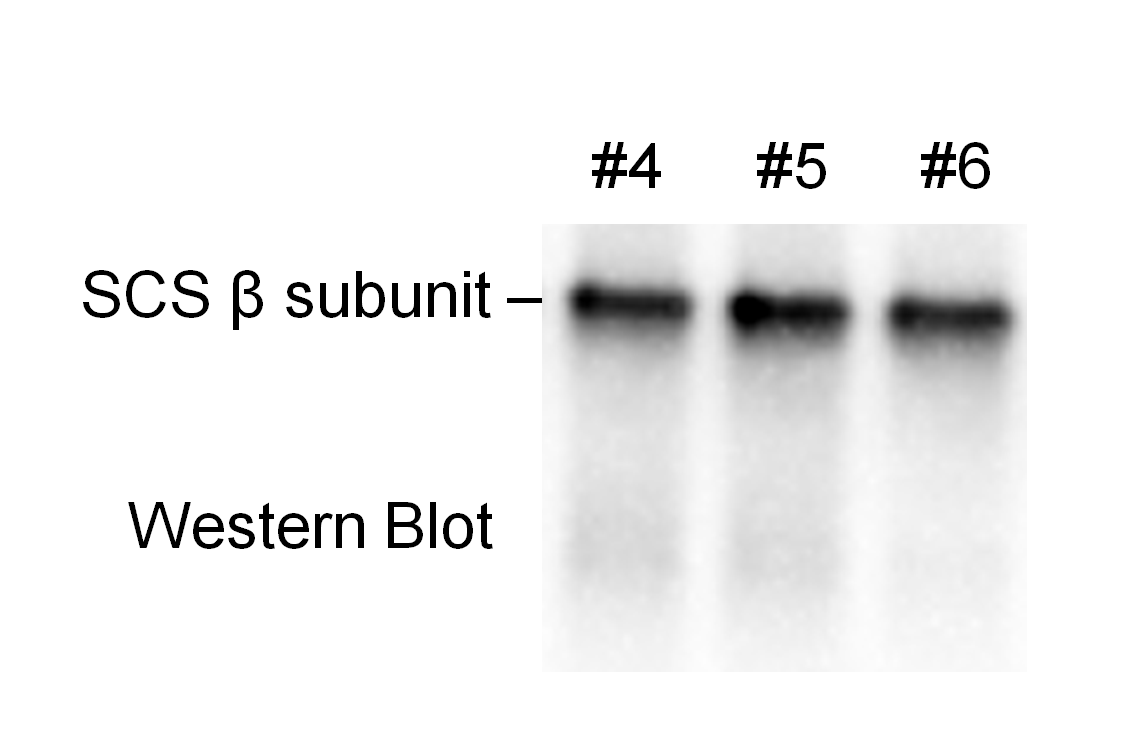

**Supplementary Figure 3**. Hyd, NuoE, NuoF of *C. churrovis* were detected in the pooled hydrogenosomal fractions by LC-MS/MS within the nanoPOTS workflow. Highlighted peptides are the combined unique peptides of each protein that were detected in three biological replicates.

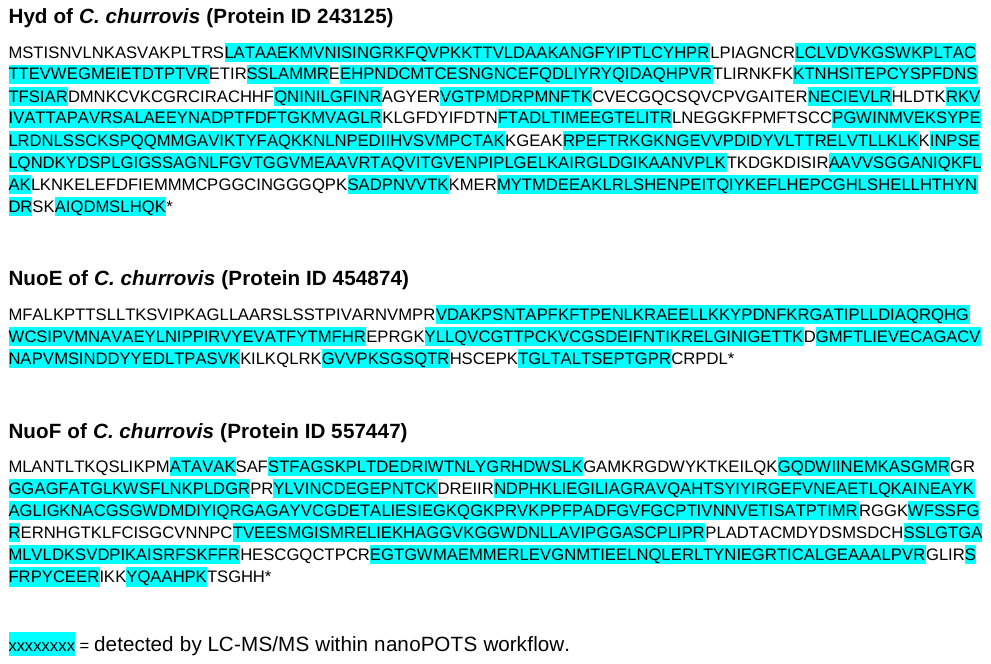

**Supplementary Figure 4.** Rank order of proteins quantified using the LC-MS/MS nanoPOTS workflow. Quantified proteins are presented as log10-transformed intensity-based absolute quantification (iBAQ) values and represent the average abundance from three biological replicates. Key proteins of interest are colored within the plot while all other are grey. ME, protein ID 462551; PFOR, protein ID 530611; Hyd, protein ID 243125; NuoE, protein ID 454874; NuoF, protein ID 557447; CcFd, protein ID 549362 in MycoCosm.

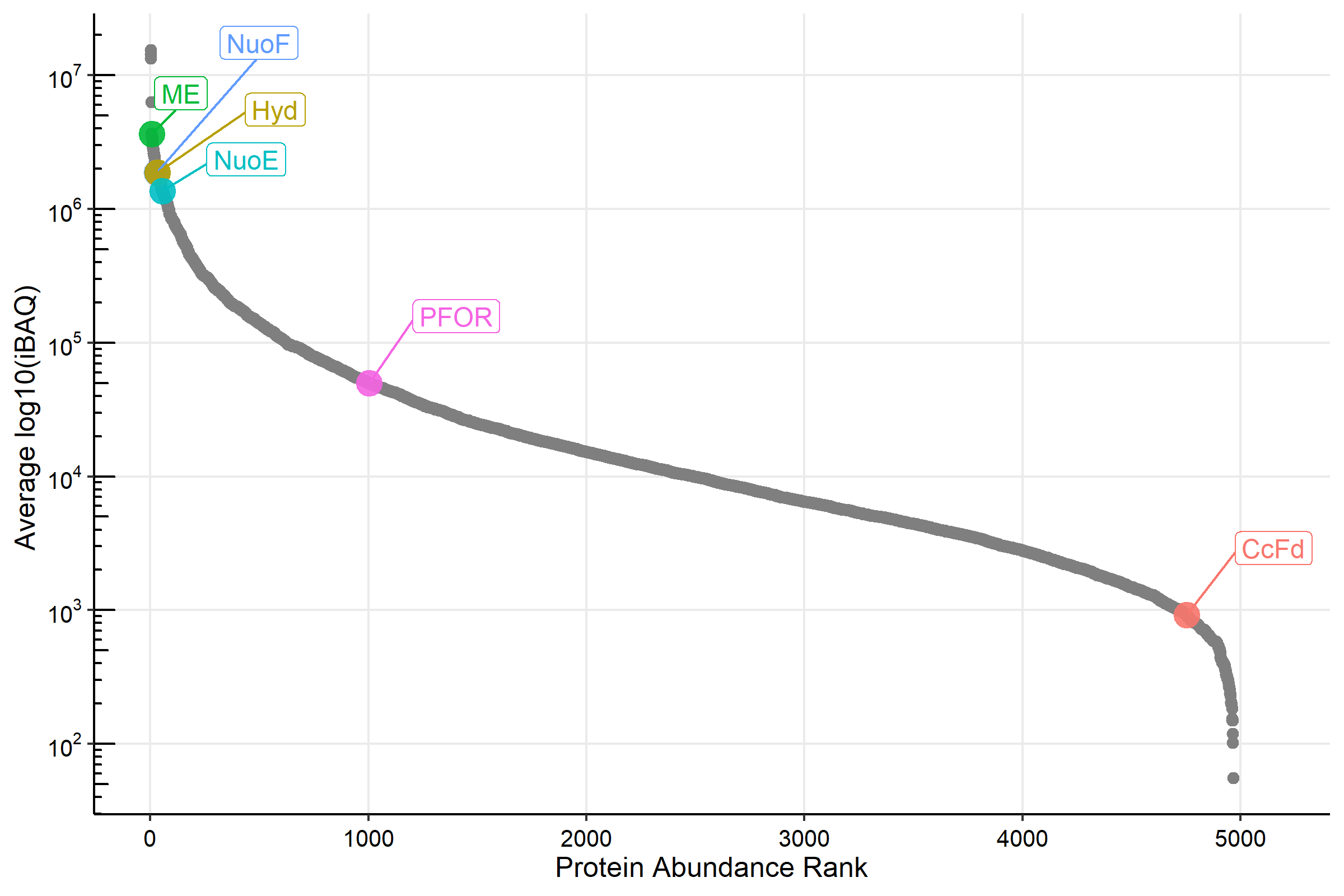

**Supplementary Figure 5**. Hyd-Strep did not reduce NAD^+^, NADP^+^, CcFd, or TvFd with H_2_ (A-D), and NuoEF-Strep did not reduce CcFd or TvFd with NADH (E-F). The assays in A-D were bubbled with H_2_ for 5 min before measuring the absorbance. CcFd or TvFd was added to the assay before starting the measurement in E-F. (A-B) NAD^+^ (1 mM) or NADP^+^ (1 mM) was not reduced with H_2_ by 5 µg Hyd-Strep in the 1-mL assay. (C-D) CcFd (30 µM) or TvFd (30 µM) was not reduced with H_2_ by about 6 µg Hyd-Strep in the 1-mL assay. The asterisk in F represents cuvette was taken out for assay mixing. (E-F) NuoEF-Strep did not reduce CcFd or TvFd (60 µM) with NADH (0.4 mM). One representative figure for each assay is shown here. Two batches of independently purified samples (Hyd-Strep, NuoEF-Strep) were tested.

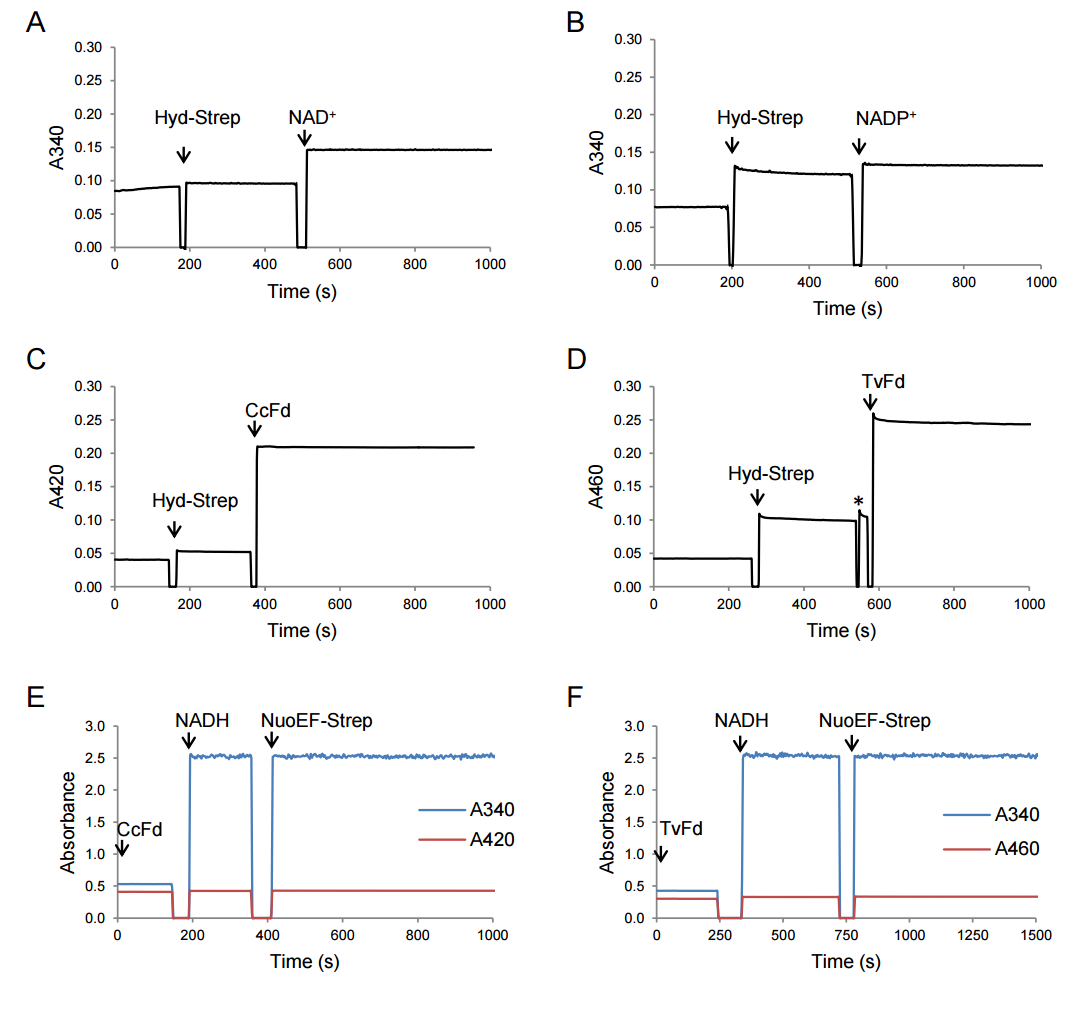

**Supplementary Figure 6**. LC-MS/MS analysis of the purified Hyd sample demonstrates the existence of Hyd-Strep in the purified Hyd samples. Similarly, NuoE and NuoF-Strep were detected in the purified NuoEF samples. Strep-tag II sequence is underlined.

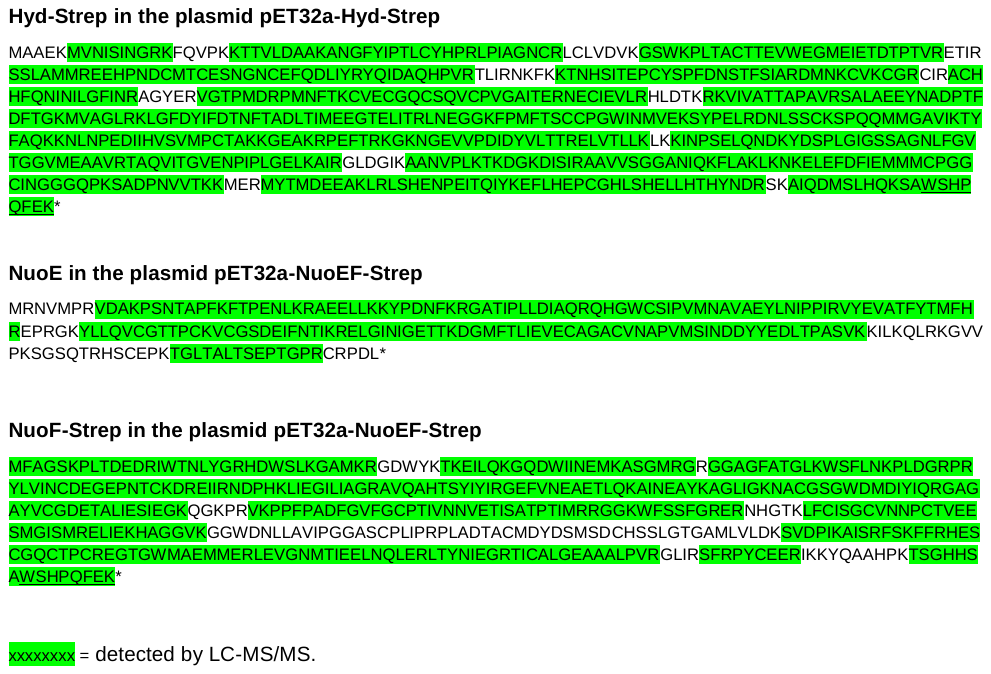

**Supplementary Figure 7**. The mixture of Hyd-Strep and NuoEF-Strep was not able to reduce NADP^+^ with H_2_ and use NADPH to form H_2_. NADP^+^, NADPH, or MV was added to the assay before starting the measurement in A-D. The assays in A and B were bubbled with H_2_ for 5 min before measuring the absorbance. (A-B) NADP^+^ was not reduced with H_2_ in the absence or presence of 30 µM CcFd in (A) or 30 µM TvFd in (B) by the mixture of 2.5 µg Hyd-Strep and 2.5 µg NuoEF-Strep in the 1-mL assay. (C) No H_2_ was formed from 2 mM NADPH by the mixture of Hyd-Strep and NuoEF-Strep before the addition of 2 mM NADH. Addition of 2 mM NADH resulted in H_2_ formation, confirming that the enzymes were functional. (D) NuoEF-Strep did not reduce MV with NADPH. Addition of 2 mM NADH lead to reduction of MV, confirming the enzyme NuoEF was functional. One representative figure for each assay is shown here. Two batches of independently purified samples (Hyd-Strep, NuoEF-Strep) were tested.

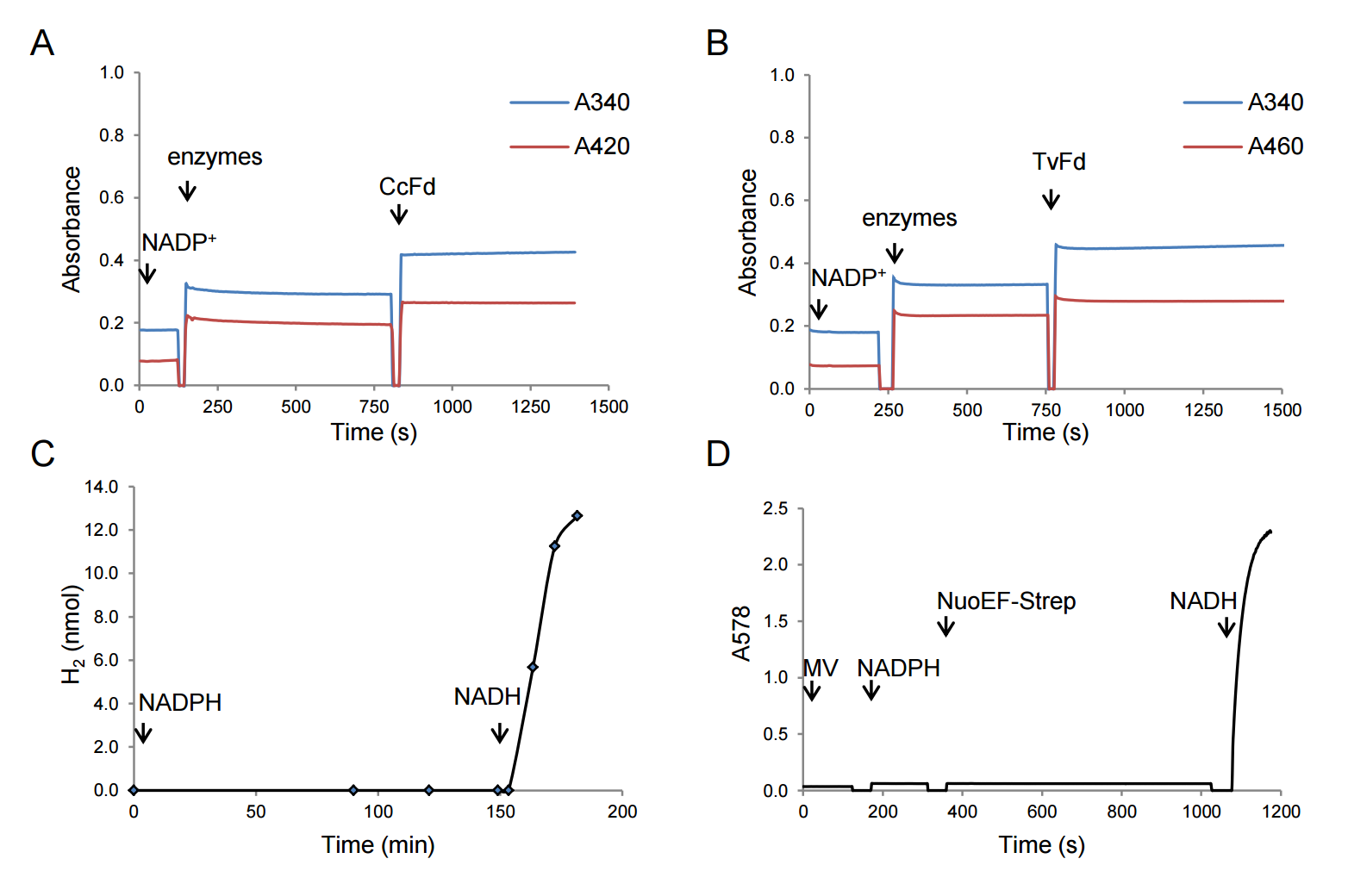

**Supplementary Figure 8**. Structure prediction of *C. churrovis* hydrogenosomal matured Hyd, NuoE, NuoF (A-C) and Hyd1ABC of *Syntrophomonas wolfei* (D-F) with ligands NAD^+^ and FAD by AlphaFold Server. Predicted 3D model with secondary structure of *C. churrovis* Hyd, NuoE, and NuoF are shown in A and C, with ipTM 0.85 and pTM 0.86. Predicted structure of *S. wolfei* Hyd1ABC are shown in D and F, with ipTM 0.88 and pTM 0.90. For A and D, the model’s color is based on the pLDTT scores, reflecting the local confidence of the model. B and E are the predicted aligned error (PAE) plots for each model. It indicates the confidence in domain orientations. For C and F, blue color represents chain Hyd or Hyd1A; green color represents chain NuoF or Hyd1B; red color represents chain NuoE or Hyd1C. (G) Predicted structures were superimposed. Color tan is for *C. churrovis* and color sky blue is for *S. wolfei*.

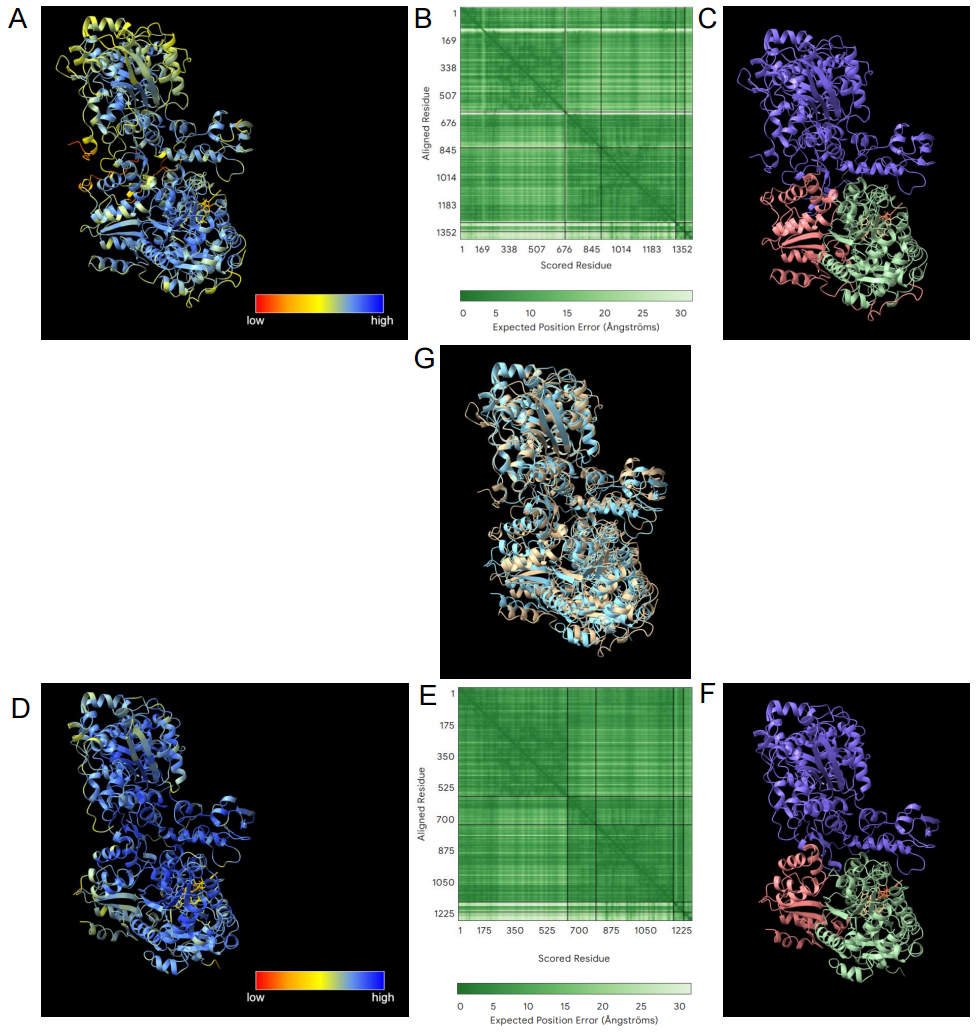

**A**
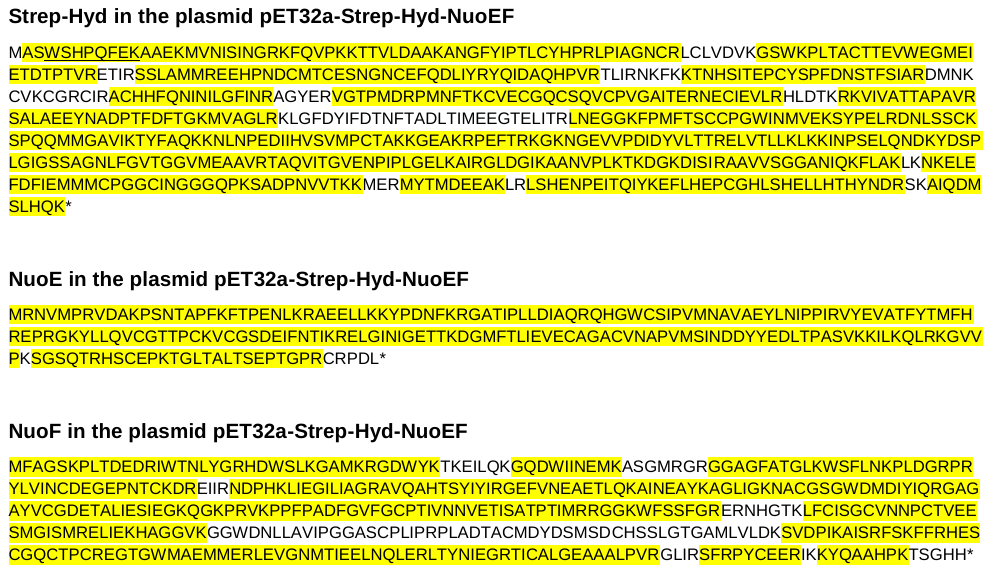

**B**
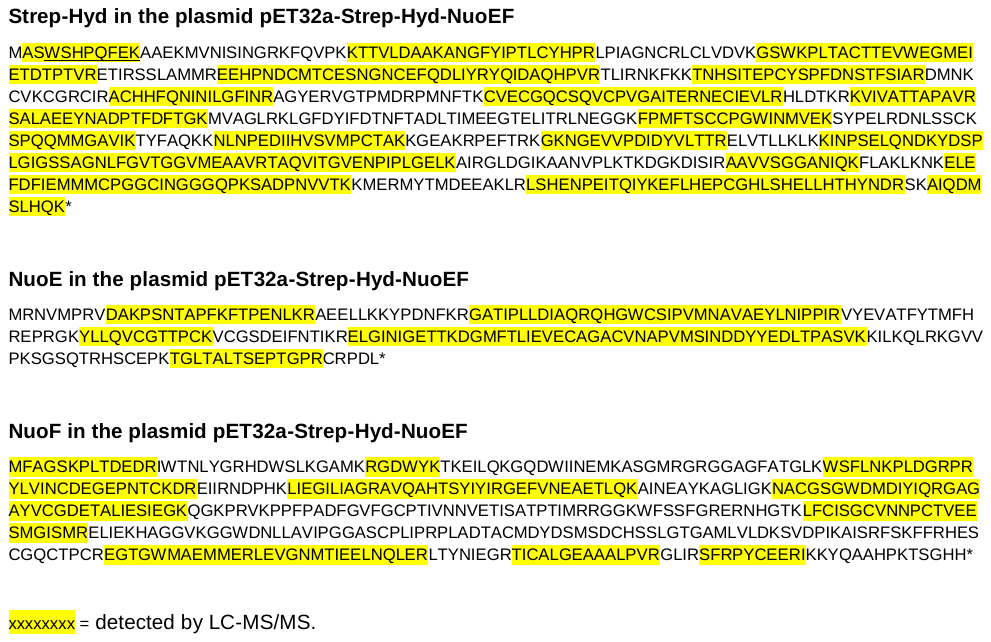

**Supplementary Figure 9**. LC-MS/MS identified peptides of the three proteins (Strep-Hyd, NuoE, NuoF) in the native PAGE gel band (A) up searing band and (B) low sharp band shown in Figure 6C. Strep-tag II sequence is underlined.

**Supplementary Figure 10**. NuoF of AF and some eukaryotes have the same key amino acids as in the experimentally verified non-bifurcating NADH-dependent enzymes. *T. maritima*, *Thermotoga maritima* ; *A. woodii*, *Acetobacterium woodii*; *S. wolfei*, *Syntrophomonas wolfei*; *S. aciditrophicus*, *Syntrophus aciditrophicus*; Anasp1, *Anaeromyces robustus* S4; Caecom1, *Caecomyces churrovis* A; Neocon1, *Neocallimastix constans* G3; NeoGFMA1, *Neocallimastix sp*. GF-Ma3-1; Neolan1, *Neocallimastix lanati*; Neosp1, *Neocallimastix californiae* G1; NeoWI3_1, *Neocallimastix sp*. WI3-B; Pecora1, *Pecoramyces sp*. F1; Pirfi3, *Piromyces finnis;* Piromy, *Piromyces sp*. UH3-1; *N. ovalis*, *Nyctotherus ovalis*; TRFO, *Tritrichomonas foetus* strain K; TVAGG3, *Trichomonas vaginalis* G3. AF NuoF IDs are also listed in Table 3. The CAA76373.1 is the Genbank accession number of *N. ovalis*. TRFO_23182 and TVAGG3_0905620 are gene IDs on TrichDB. SLBB domain means soluble-ligand-binding-beta-grasp domain. The numbers before peptide sequences represent positions of the first amino acid in each column.

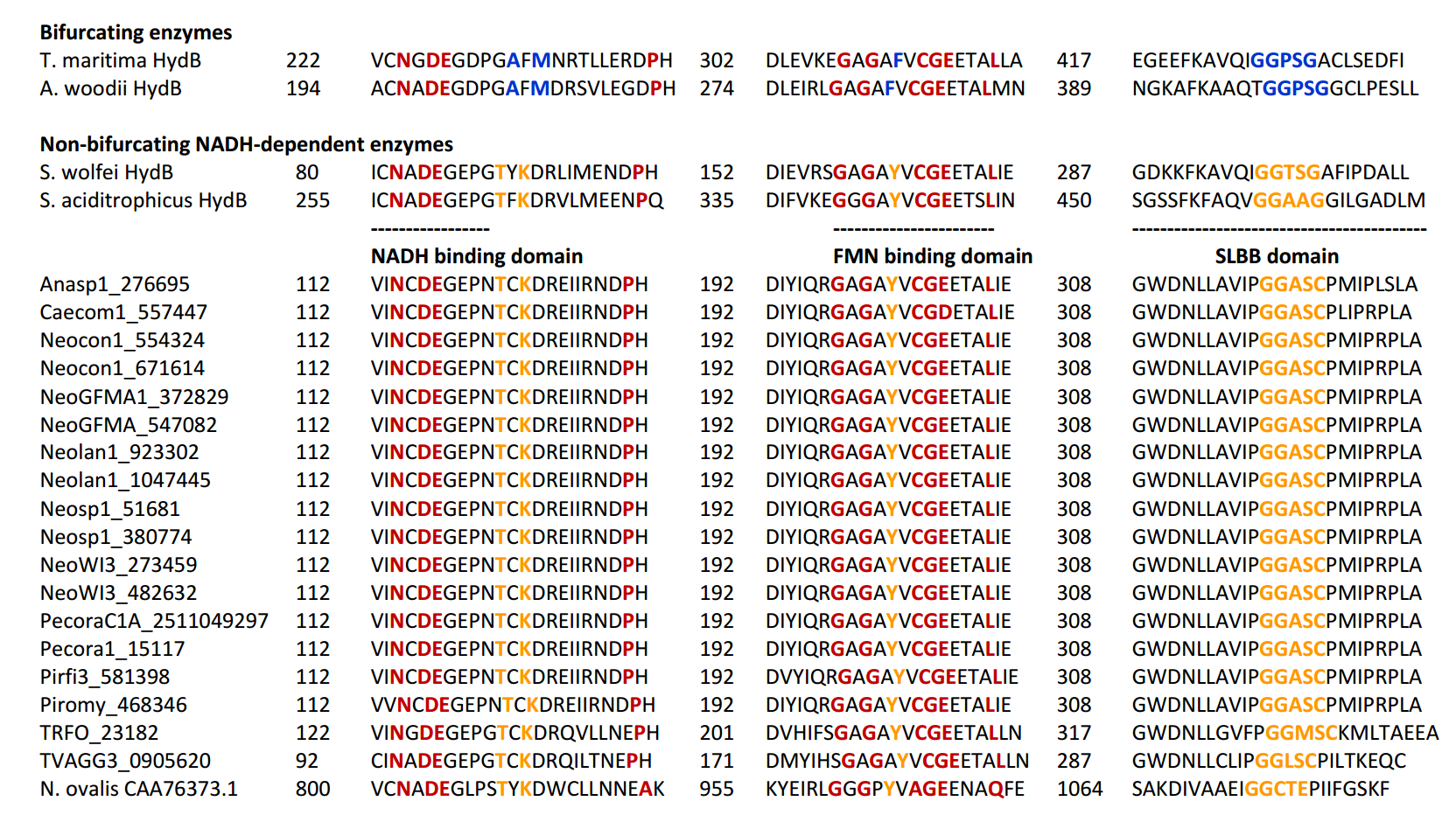
